# Genetic Dissociation of Circadian Prediction and Behavioral Output by a Calbindin1+ Dopamine Neuron Population

**DOI:** 10.64898/2026.03.27.714759

**Authors:** Andrew P. Villa, Jacqueline Trzeciak, Damien Wolfe, David E. Ehichioya, Jeffrey Falkenstein, Justine T. Wong, Renqi Wu, Lauren Dimalanta, Wyatt Kaiban, Jaskaran Dhanoa, Gregory Stevens, Moueez Shah, Fernando Garcia, Lori Scarpa, Chris Chalfoun, Larry S. Zweifel, Rajeshwar Awatramani, Richard D. Palmiter, Martin Darvas, Shin Yamazaki, Andrew D. Steele

**Affiliations:** Department of Biological Sciences, California State Polytechnic University Pomona; Pomona, CA; USA; Department of Neuroscience; Peter O’Donnell Jr. Brain Institute, UT Southwestern Medical Center, Dallas, TX; USA; Division of Biology and Bioengineering, California Institute of Technology, Pasadena, CA; USA; Department of Psychiatry and Behavioral Sciences, Department of Pharmacology, University of Washington, Seattle, United States; Department of Neurology, Northwestern University, Chicago, IL 60611, USA; Howard Hughes Medical Institute and Departments of Biochemistry and Genome Sciences, University of Washington, Seattle, WA, USA; Department of Laboratory Medicine and Pathology, University of Washington, Seattle, WA, USA

## Abstract

Anticipating daily food availability is a conserved circadian behavior that persists even in animals lacking the suprachiasmatic nucleus, yet its neural substrates remain poorly defined. Previous studies implicated dopamine signaling in food anticipatory activity (FAA) but lacked the resolution to identify the responsible neuronal population. Here, we conditionally deleted *tyrosine hydroxylase* (*Th*) from molecularly defined dopamine neuron populations in mice. Broad deletion of *Th* in dopamine transporter-expressing neurons nearly abolished FAA, whereas restoration of *Th* in substantia nigra dopamine neurons rescued anticipatory locomotion. Surprisingly, deletion of *Th* from several large dopamine neuron populations had little effect on FAA. In contrast, deletion using *Calb1^Cre^*, targeting only ∼25% of substantia nigra dopamine neurons, produced a profound FAA deficit. Notably, these mice exhibited a profound loss of anticipatory locomotion while retaining substantial anticipatory food-seeking behavior. These findings identify a small Calbindin1+ dopamine population required for anticipatory locomotion and demonstrate that distinct behavioral components of food anticipation can be genetically dissociated.

## Introduction

Circadian rhythms in physiology and behavior are governed by cell-autonomous transcriptional-translational feedback loops operating in nearly all cells.^1,2^ In the mammalian brain, the suprachiasmatic nucleus (SCN) functions as the central circadian pacemaker, aligning internal rhythms with the environmental light-dark cycle.^1^ Because most animals feed in phase with their active time, the role of feeding as an entraining signal is often overlooked. However, when rodents are fed during their rest phase, physiological rhythms such as body temperature and corticosterone secretion, along with a significant increase in locomotor activity, shift to precede mealtime.^3^ This food anticipatory activity (FAA) persists even in animals with SCN lesions, suggesting that the SCN plays little or no role in regulating food-entrained rhythms.^4^ Furthermore, disrupting the circadian rhythm in the SCN with a triple deletion of *Period 1*, *2*, and *3* did not impair nose poking in anticipation of scheduled feeding in mice and suggested that an SCN-independent light-entrained clock may be involved in predicting circadian feeding time.^5^

The search for brain regions that respond to food in a circadian fashion—analogous to how the SCN responds to light—has been long and inconclusive.^6–8^ The absence of a clearly defined food-entrainable oscillator (FEO), combined with evidence that rodents can anticipate multiple meals, has led to the hypothesis that FAA is governed by a distributed network of FEOs.^3,6,9,10^ Supporting this view, manipulations targeting diverse brain regions and circuits impair but do not eliminate FAA. These include cerebellar dysfunction, disrupted connectivity between the dorsomedial hypothalamus and SCN, manipulation of hunger-sensitive hypothalamic neurons, and genetic or pharmacological disruptions of dopamine signaling.^11–18^

There is growing interest in how dopamine regulates biological rhythms, particularly FAA.^16,19,20^ Because dopamine is essential for feeding, locomotor activity, habit formation, and most reward-directed behaviors,^21,22^ dopamine neurons may serve as FEOs themselves or as critical regulators of their motor output. In rodents, dopamine tone in the dorsal striatum (DS) fluctuates diurnally, with peak levels at night and troughs during the day, as measured by in vivo microdialysis and voltammetry.^23^ This rhythm is driven by the dopamine transporter (DAT, encoded by *Slc6a3*), whose expression cycles in a circadian manner.^17^ Notably, DAT-knockout (KO) mice lack this dopamine rhythm.^23^ Moreover, core clock gene expression (e.g., *Period 2*) in DS neurons is rhythmic only when dopamine innervation is intact.^24^ These findings suggest that dopamine plays a key role in synchronizing striatal circadian function, although direct evidence that dopamine levels in the DS are modulated by restricted feeding is still lacking.^16^

In prior studies, we demonstrated that dopamine signaling via the dopamine D1 receptor (D1R) is important for FAA.^17^ Mice lacking D1R exhibit normal rises in pre-prandial core body temperature but have severely reduced FAA in response to scheduled, calorie-restricted (CR) feeding. Although follow-up studies showed that the FAA deficit in D1R KO mice was less severe than initially reported, all four tested lines displayed significant impairment.^25,26^ Other receptor subtypes may also contribute: dopamine D2 receptor (D2R) activation can advance FAA onset in rats, and inhibition of either D1R or D2R signaling reduces FAA.^15,27^ However, these pharmacological studies rely on systemic drug delivery, which lacks anatomical precision and is confounded by presynaptic D2 auto-receptor effects.

Genetic studies restoring dopamine selectively to the DS in otherwise dopamine-deficient mice suggest that FAA is mediated through nigrostriatal dopamine pathways.^17^ Using *paired-like homeodomain 3* (*Pitx3*) mutant mice, we demonstrated that a minimal nigrostriatal pathway is sufficient to support FAA; however, follow-up studies in this model were hindered by pleiotropic phenotypes present in these hypomorphic mutants (e.g., retinal dysfunction and metabolic abnormalities).^28^ Based on these results, we sought to leverage the molecular heterogeneity of dopamine neurons and conditional genetic tools to identify a dopamine subpopulation projecting to the DS that is required for FAA. Single-cell PCR and RNA-seq analyses have revealed multiple molecularly distinct subtypes of midbrain dopamine neurons, each with unique physiological and anatomical properties.^29–31^ These subtypes can be selectively targeted using Cre-lox-based genetic strategies to dissect their functional roles in FAA. To this end, we performed a series of conditional deletions of *tyrosine hydroxylase* (*Th*), the rate-limiting enzyme for catecholamine biosynthesis, to screen for genetically defined dopamine populations required for FAA. Through this approach, we identified several major dopamine neuron populations that are dispensable for FAA and a small Calbindin1-lineage population that is selectively required for anticipatory locomotion. These findings provide new insight into the organization of dopaminergic circuits underlying food anticipation.

## Results

### Conditional Deletion of Tyrosine Hydroxylase in Slc6a3-Expressing Neurons Severely Impairs Food Anticipatory Activity

To identify dopamine neurons required for FAA, we conditionally deleted *Th* using *Pitx3^Cre^*.^32^ Consistent with prior work, this manipulation resulted in perinatal lethality, reflecting near-complete dopamine deficiency (p < 0.0001, Log-Rank (Mantel-Cox) test; **Figure S1A**).^21^ We therefore turned to *Slc6a3^Cre^* (DAT-Cre) mice to selectively eliminate *Th* in DAT-expressing neurons while preserving viability thereby targeting the major population of midbrain dopaminergic neurons.^33–35^

Double labeling for TH and DAT in wild-type mice revealed a small population of TH-positive, DAT-negative neurons in the substantia nigra (SN) and ventral tegmental area (VTA) in the midbrain, indicating that *Slc6a3^Cre^*-mediated recombination would not eliminate all dopaminergic neurons (**Figure 1A-B**; **S1B**). Consistent with this, *Slc6a3^Cre^;Th^Flox^* conditional knockout (DAT-TH-cKO) mice exhibited a robust loss of TH immunoreactivity throughout DAT-expressing regions of the midbrain (consistent with efficient recombination in DAT-lineage neurons, p = 0.0001; unpaired two-tailed t-test with Welch’s Correction; **Figure 1C**). A sparse residual population of TH-positive neurons was revealed using whole-brain CLARITY-based immunolabeling^36^ (**Figure S1C; Video S1-5**), consistent with DAT-negative dopaminergic neurons observed in wild-type mice by both TH/DAT immunostaining **(Figure 1A)** and *Th/Slc6a3 in situ* hybridization (**Figure S1B**). This model allowed us to test whether DAT-expressing dopaminergic neurons are required for FAA.

**Figure 1.**
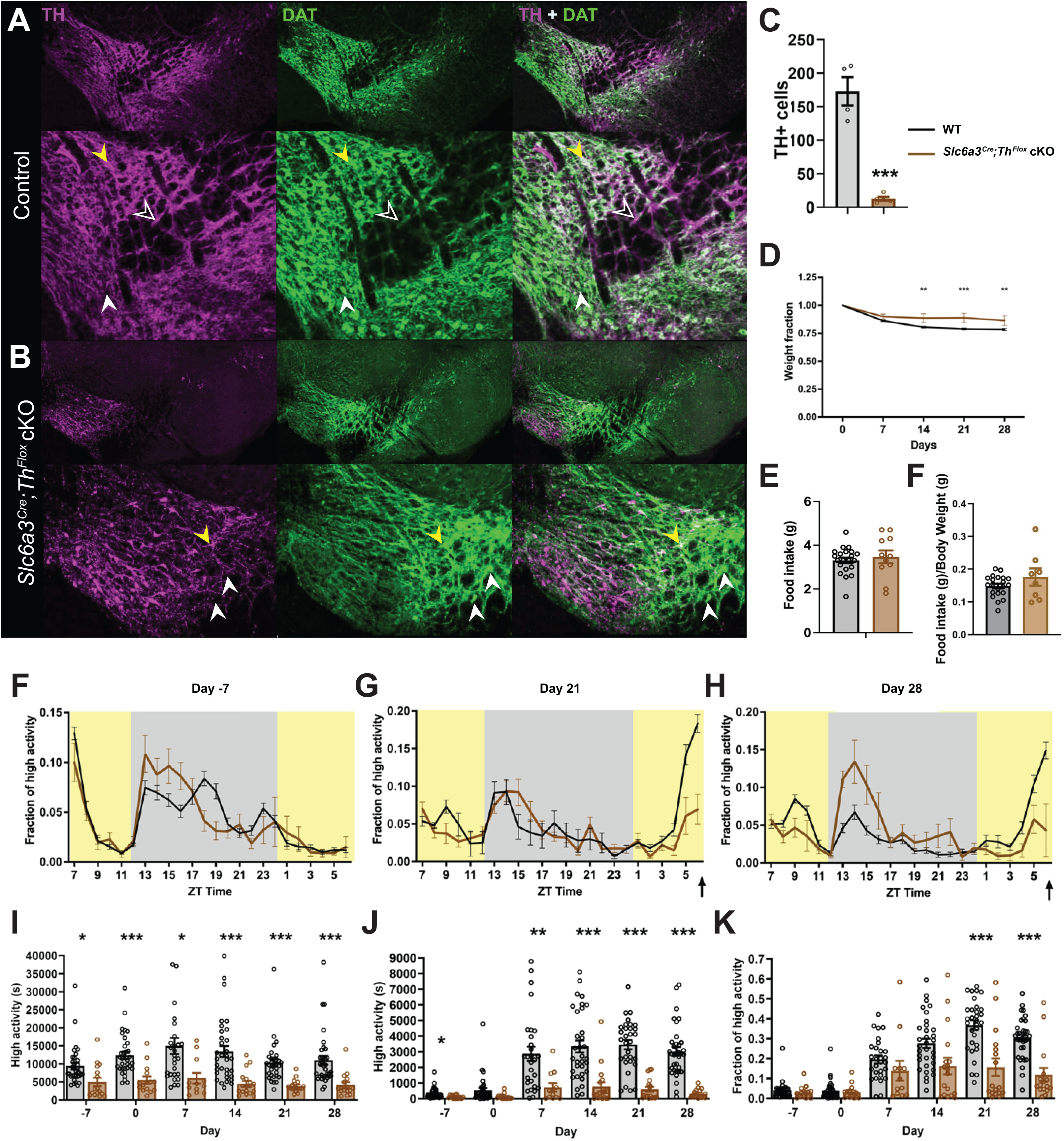
Conditional Deletion of *Tyrosine Hydroxylase* in *Slc6a3^Cre^* Neurons Severely Impairs FAA. A) Confocal imaging of immunofluorescence antibody staining of TH (magenta) and DAT (green) in control midbrain. and *Slc6a3^Cre^;Th^Flox^* cKO. Overlap (far right) shows co-expression (white) of these markers. Yellow arrowhead indicates TH-DAT double positive cell; open arrowheads indicate TH^+^/DAT^-^ cells; while white arrowhead indicates a TH^-^/DAT^+^ cell. B) Confocal imaging of immunofluorescence antibody staining of TH and DAT in *Slc6a3^Cre^;Th^Flox^* cKO midbrain. The yellow arrowhead indicates a TH-DAT double positive cell that escaped Cre-mediated deletion; while the white arrowheads indicate TH-/DAT+ cell likely from Cre-mediated deletion. C) Cell count of TH^+^ cells in WT (black) and *Slc6a3^Cre^;Th^Flox^* cKO (brown). D) Normalized body weight of WT and *Slc6a3^Cre^;Th^Flox^* cKO mice over the course of the study; weight was normalized by dividing the weekly weight by the Day 0 weight, which was taken prior to beginning CR. E) Raw food intake of WT and *Slc6a3^Cre^;Th^Flox^* cKO mice while on AL. F) Normalized food intake of WT and *Slc6a3^Cre^;Th^Flox^* cKO mice while on AL; data is normalized by dividing total AL raw food intake by initial body weight. G-I) Normalized high activity on days -7 (G), 21 (H), and 28 (I); data is normalized by dividing the seconds of high activity in the premeal window by the total daily high activity. J) Total daily high activity in seconds throughout the course of the study. K) High activity in seconds during the 3-hour premeal window throughout the course of the study. L) Normalized high activity during the 3-hour premeal window; data is normalized by dividing the seconds of high activity in the premeal window by the total daily high activity. Data are mean +/- SEM. Statistical significance of behavior was determined using a Mixed Effect’s Analysis with Sidak’s Multiple Comparisons Test. Statistical significance of cell counts, feeding and weight were determined using an Unpaired Two-tailed t-test with Welch’s Correction. ∗p < 0.05, ∗∗ p < 0.01, *** p < 0.001.

Phenotypically, *Slc6a3^Cre^;Th^Flox^* cKO mice exhibited reduced postnatal viability, and many died without intervention. Providing moistened food on the cage floor improved this viability. When fed *ad libitum* (AL), DAT-TH-cKO mice consumed comparable amounts to controls (**Figure 1E),** which persisted when normalized to body weight (3.8 ± 0.4 g vs. 3.3 ± 0.1 g, p = 0.363, unpaired two-tailed t-test with Welch’s Correction; **Figure 1F**). To assess FAA, mice were restricted to 60–70% of AL intake, delivered daily at ZT7 for 28 days. Initial body weight measurements over the course of the CR period showed WT and DAT-TH-cKO groups to be fairly similar (**Figure S1D).** However, normalizing body weights taken during this CR period indicates cKO mice are more resistant to weight loss compared to WT counterparts after day 7 of study (day 14, p = 0.0029; day 21, p = 0.0001; day 28, p = 0.0028; mixed effects analysis with Sidak’s multiple comparisons test; **Figure 1D**).

We quantified home-cage activity using computer vision-based tracking software, which measured high-intensity behaviors (walking, jumping, hanging, rearing) across the circadian cycle.^37^ Baseline-activity patterns were assessed prior to timed CR feeding on “day -7”, during which time both control and cKO mice exhibited comparable levels of normalized activity (**Figure 1G**). However, by days 21 and 28 of timed CR feeding, the DAT-TH-cKO mice demonstrated only a small increase in premeal high activity behaviors compared to controls (**Figure 1H-I**). As expected for a dopamine-deficient model, total high activity levels were reduced in DAT-TH-cKO mice compared to controls for days 0, 7, 14, and 28 (day -7, p = 0.0066; day 0, p = 0.0008; day 7, p < 0.0001; day 14, p = 0.0090; day 28, p = 0.0050; mixed effects analysis with Sidak’s multiple comparisons test; **Figure 1J).** By day 7 of CR feeding, control mice showed marked increases in pre-meal activity, whereas DAT-TH-cKO mice exhibited only minimal changes (p _≤_ 0.0001; mixed effects analysis with Sidak’s multiple comparisons test; **Figure 1K**).

To account for baseline hypoactivity, we calculated normalized high activity, defined as the fraction of total daily activity occurring in each hour.^7,17,38^ This normalization controls for inter-individual differences in overall activity and allows comparisons across genotypes and experiments. Using this metric, DAT-TH-cKO mice exhibited a marked impairment in FAA. Control mice redistributed up to 30% of their daily activity to the 3 hours preceding feeding. DAT-TH-cKO mice exhibited markedly reduced redistribution of activity to the pre-meal period, with values remaining near baseline across animals (p < 0.001; Mixed effects analysis with Sidak’s multiple comparisons test; **Figure 1L**). Thus, dopamine synthesis in DAT-expressing neurons is required for the normal expression of FAA.

### Viral Restoration of TH in the SN Restores FAA in Slc6a3^Cre^;Th^Flox^ cKO mice

To test whether restoring dopamine synthesis in DAT-expressing neurons within the SN could rescue FAA, DAT-TH-cKO mice were subjected to bilateral injections of a Cre-dependent AAV expressing TH tagged with a red fluorescent protein (RFP-TH) marker (**Figure 2A-B**). To verify restored TH expression in the SN, we performed immunohistochemistry for TH and either RFP (DAT-TH-cKO rescue) or GFP (DAT-TH-cKO negative control). Mice with _≥_50 RFP+/TH+ neurons per SN were classified as successfully rescued. The viral rescue was partially successful as we observed modest restoration of TH expression in SN neurons (**Figure 2A; S2A-B**). Quantification of AAV-DIO-TH-P2A-RFP–injected DAT-TH-cKO rescue mice showed a significantly greater number of TH+ neurons (50 ± 10 cells/section averaged per animal, n = 8 mice; **Figure S2C**), comparably above the number of GFP+ cells in AAV9-DIO-GFP DAT-TH-cKO controls (17 ± 2 cells/section, n = 6 mice; **S2D).** When comparing the quantification of TH+ neurons in both groups, RFP-TH cKO rescue mice exhibited a significantly larger number of TH+ neurons (p = 0.017, unpaired two-tailed t-test with Welch’s Correction; **Figure 2C**), confirming successful viral rescue of TH expression. Notably, TH restoration failed to alter body weight (**Figure S2E),** even when normalized (p = 0.981; mixed effects analysis with Sidak’s multiple comparisons test; **Figure 2D)**, but it did initially increase food intake during AL feeding in TH-RFP rescue mice (p = 0.0162; unpaired two-tailed t-test with Welch’s Correction; **Figure S2F)**. However, total AL food intake remained comparable between groups when normalized to body weight (**Figure 2E**). To assess FAA, mice were restricted to 60–70% of AL intake, delivered daily at ZT7 for 28 days. Overall locomotor activity during the CR period was unaffected (p > 0.05; Mixed effects analysis with Sidak’s multiple comparisons test; **Figure 2I**).

**Figure 2.**
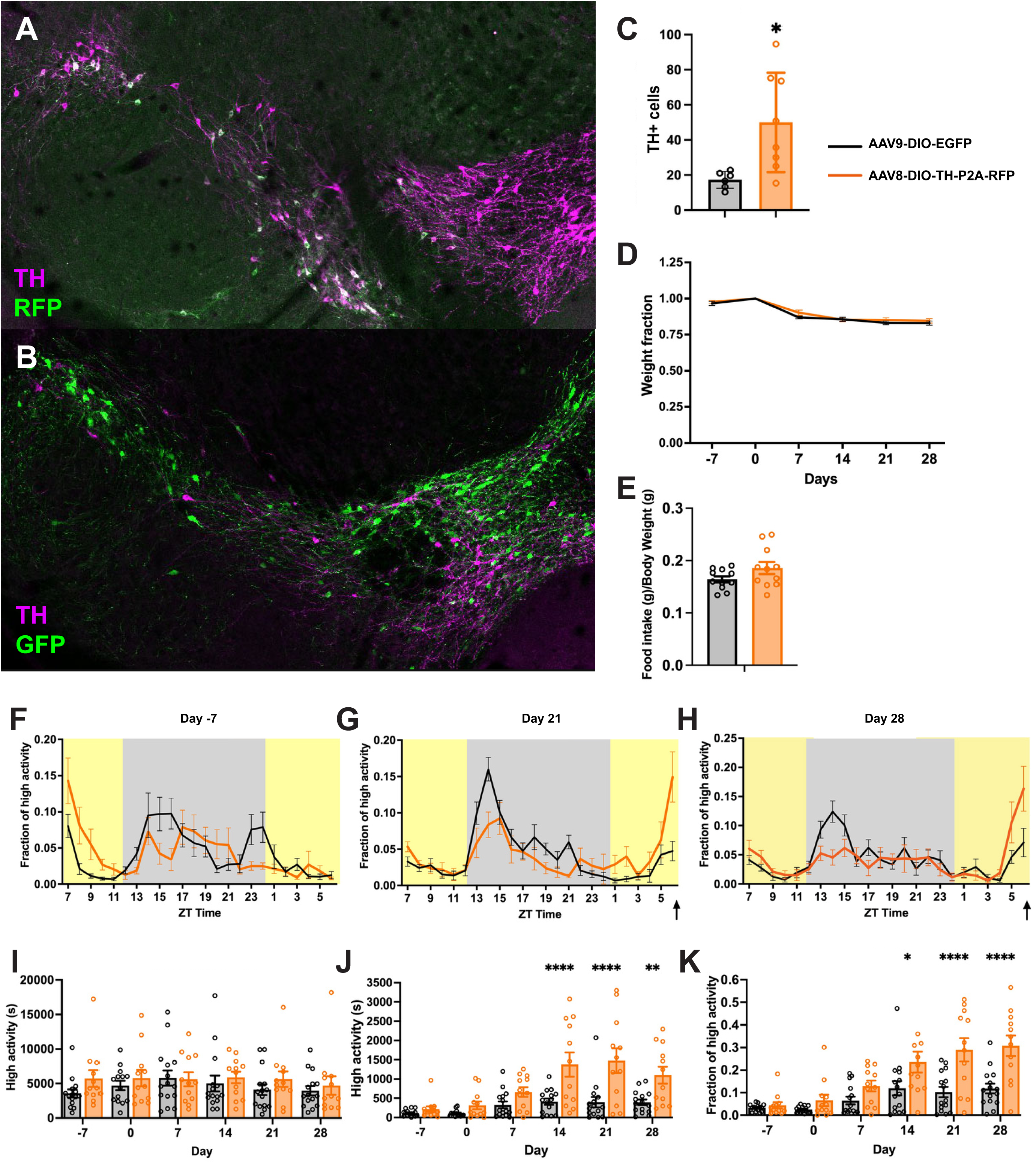
Viral rescue of TH expression in the SN restores FAA in Slc6a3*^Cre^;Th^Flox^* mice. A) Confocal imaging of immunofluorescence antibody staining of TH (magenta) and RFP (green) limited viral rescue of TH depleted neurons in *Slc6a3^Cre^;Th^Flox^* cKO mice. Overlap shows the co-expression (white) of these markers in cKO mice injected with AAV-DIO-TH-P2A-RFP into the SN. B) Confocal imaging of immunofluorescence antibody staining of TH (magenta) and GFP (green) negative control neurons in *Slc6a3^Cre^;Th^Flox^* cKO mice. Overlap shows the co-expression (white) of these markers in cKO mice injected with AAV-DIO-EGFP into the SN. C) Comparison of Midbrain SN TH+ cell counts per brain section in AAV-GFP (black) control mice and AAV-RFP-TH (orange) rescue mice. D) Normalized body weight of TH rescue and negative control *Slc6a3^Cre^;Th^Flox^* cKO mice over the course of the study; weight was normalized by dividing the weekly weight by the Day 0 weight, which was taken prior to beginning CR. E) Normalized food intake of TH rescue and negative control *Slc6a3^Cre^;Th^Flox^* cKO mice while on AL; data is normalized by dividing total AL raw food intake by initial body weight. F-H) Normalized high activity on days –7 (F), 21 (G), and 28 (H); data is normalized by dividing the seconds of high activity in the premeal window by the total daily high activity. I) Total daily high activity in seconds throughout the course of the study. J) High activity in seconds during the 3-hour premeal window throughout the course of the study. K) Normalized high activity during the 3-hour premeal window; data is normalized by dividing the seconds of high activity in the premeal window by the total daily high activity. Data are mean +/- SEM. Statistical significance of behavior was determined using Sidak’s Multiple Comparisons Test. Statistical significance of cell counts, feeding and weight were determined using unpaired Welch’s T test. ∗p < 0.05, ∗∗ p < 0.01, *** p < 0.001

Collectively, these data indicate that any behavioral rescue was not due to global improvements in feeding or locomotor activity.

Despite persistent hypoactivity, AAV-mediated TH restoration in SN robustly rescued FAA. Rescued mice developed significant pre-meal increases in activity that emerged by day 14 of CR and persisted through day 28 **(Figure 2F-H**), in contrast to GFP-injected controls, which failed to develop pre-meal activity (Day -7, Day 0, and Day 7, p >0.05; Day 14, p < 0.0001; Day 21, p < 0.0001; Day 28, p = 0.0071; mixed effects analysis with Sidak’s multiple comparisons test; **Figure 2J-K).** Thus, dopamine synthesis in a surprisingly small number of SN neurons is sufficient to restore anticipatory locomotion (50 ± 10 cells/section averaged per animal, n = 8 mice; **Figure S2C**).

### Conditional Deletion of Tyrosine Hydroxylase in Multiple Molecularly Defined Dopamine Neuron Populations Minimally Affects FAA

Because FAA was nearly abolished in DAT-TH-cKO mice yet restored by re-expression of TH in a limited number of SN neurons, we next asked whether specific molecularly defined dopamine neuron populations are required for FAA. We therefore performed a systematic conditional knockout screen across five genetically distinct dopamine neuron populations previously identified through molecular profiling studies.^29^ To delete *Th* in distinct subsets of midbrain dopamine neurons, we employed five *Cre* driver lines: *Crhr1^Cre^*,^39^ *Foxp2^Cre^*,^40^ *Ntsr1^Cre^*,^41^ *Sox6^Cre^*,^42^ and *Slc17a6^Cre^*.^43^ In each line, TH deletion substantially reduced the number of TH^+^ neurons in the SN. Although FAA developed slightly more slowly in two lines, all five models ultimately achieved comparable levels of FAA.

In *Crhr1^Cre^;Th^Flox^* cKO (Crhr1-TH-cKO) mice, TH expression in the ventral midbrain was markedly reduced (**Figure 3A**). The number of TH^+^ neurons in the SN per section decreased from 129 ± 15 in controls to 23 ± 2 in cKOs (p = 0.0019, unpaired two-tailed t-test with Welch’s Correction; **Figure 3B**). Despite this substantial loss of TH, FAA developed normally in Crhr1-TH-cKO mice (p > 0.05; mixed effects analysis with Sidak’s multiple comparisons test; **Figure 3C**). Total locomotor activity was modestly reduced at several time points (Day –7, p = 0.163; Day 0, p = 0.006; Day 7, p = 0.418; Day 14, p = 0.012; Day 21, p = 0.072; Day 28, p = 0.089; mixed effects analysis with Sidak’s multiple comparisons test; **Figure S3A**), but raw premeal activity did not differ significantly between controls and cKO mice (p > 0.05; mixed effects analysis with Sidak’s multiple comparisons test; **Figure S3B**). Crhr1-TH-cKO mice were smaller than littermate controls throughout the study (Day –7, p = 0.042; Day 0, p = 0.001; Day 7, p = 0.016; Day 14, p = 0.037; Day 21, p = 0.046; Day 28, p = 0.0317; mixed effects analysis with Sidak’s multiple comparisons test; **Figure S3C**). However, normalized body weight during CR was largely similar between genotypes (Day –7, p = 0.007; Day 0, p > 0.999; Day 7, p = 0.686; Day 14, p = 0.299; Day 21, p = 0.207; Day 28, p = 0.539; mixed effects analysis with Sidak’s multiple comparisons test; **Figure S3D**). Food intake was also similar between groups (p > 0.05; unpaired two-tailed t-test with Welch’s Correction, **Figure S3E**), although body weight–normalized intake showed a mild increase in cKO mice (p = 0.0189; unpaired two-tailed t-test with Welch’s Correction; **Figure S3F**).

**Figure 3.**
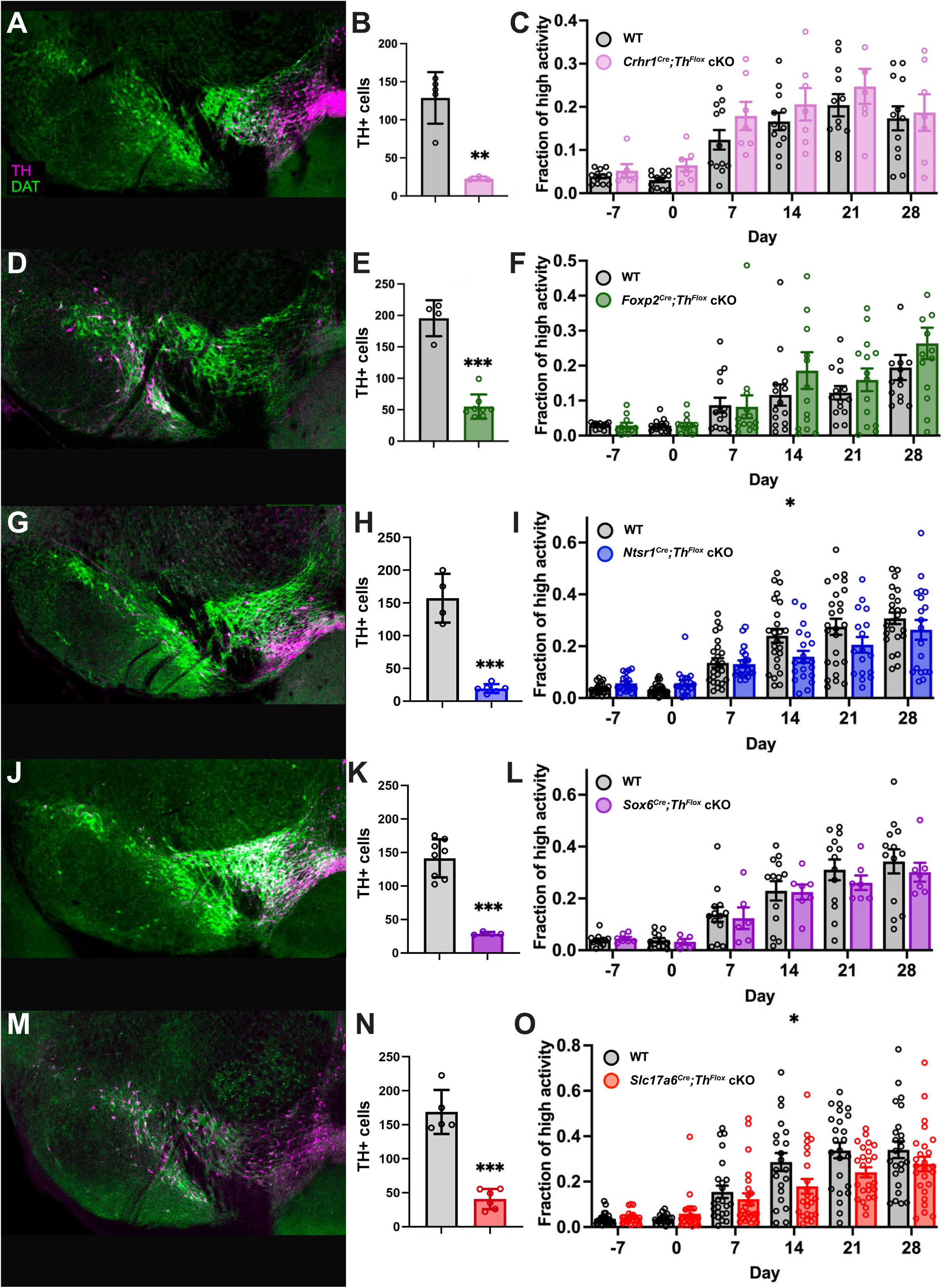
Conditional Deletion of Tyrosine Hydroxylase in *Crhr1-, Foxp2-, Ntsr1-, Sox6-, and Slc17a6^Cre^* Neurons Minimally Alters FAA. A) Confocal imaging of immunofluorescence antibody staining of TH (magenta) and DAT (green) in the midbrain of *Crhr1^Cre^;Th^Flox^* cKO brain tissue. B) Quantification of TH staining in the SN of WT (gray) and *Crhr1^Cre^;Th^Flox^* cKO (pink) midbrain tissue. C) Normalized high activity of control and *Crhr1^Cre^;Th^Flox^* cKO mice during the 3-hour premeal window over the course of the study. D) Confocal imaging of immunofluorescence antibody staining of TH (magenta) and DAT (green) in the midbrain of *Foxp2^Cre^;Th^Flox^* cKO and control brain tissue. E) Quantification of TH staining in the SN of WT (grey) and *Foxp2^Cre^;Th^Flox^* cKO (green) midbrain tissue. F) Normalized high activity of *Foxp2^Cre^;Th^Flox^* WT and cKO mice during the 3-hour premeal window over the course of the study. G) Confocal imaging of immunofluorescence antibody staining of TH (magenta) and DAT (green) in the midbrain of *Ntsr1^Cre^;Th^Flox^* cKO brain tissue. H) Quantification of TH staining in the SN of WT (grey) and *Ntsr1^Cre^;Th^Flox^* cKO (blue) midbrain tissue. I) Normalized high activity of *Ntsr1^Cre^;Th^Flox^* WT and cKO mice during the 3-hour premeal window over the course of the study. J) Confocal imaging of immunofluorescence antibody staining of TH (magenta) and DAT (green) in the midbrain of *Sox6^Cre^;Th^Flox^* cKO brain tissue. K) Quantification of TH staining in the SN of WT (grey) and *Sox6^Cre^;Th^Flox^* cKO (purple) midbrain tissue. L) Normalized high activity of *Sox6^Cre^;Th^Flox^* WT and cKO mice during the 3-hour premeal window over the course of the study. M) Confocal imaging of immunofluorescence antibody staining of TH (magenta) and DAT (green) in the midbrain of *Slc17a6^Cre^;Th^Flox^* cKO brain tissue. N) Quantification of TH staining in the SN of WT (grey) and *Slc17a6^Cre^;Th^Flox^* cKO (red) midbrain tissue. O) Normalized high activity of *Slc17a6^Cre^;Th^Flox^* WT and cKO mice during the 3-hour premeal window over the course of the study. Data are mean +/- SEM. Statistical significance of behavior was determined using Sidak’s Multiple Comparisons Test, statistical significance of cell counts was determined using unpaired Welch’s T test. ∗p < 0.05, ∗∗ p < 0.01, *** p < 0.001.

In *Foxp2^Cre^;Th^Flox^* cKO (Foxp2-TH-cKO) mice, TH expression was also strongly reduced in the ventral midbrain (**Figure 3D**). The number of TH+ SN neurons decreased from 196 ± 29 in controls to 55 ± 19 in cKOs (p = 0.0006; unpaired two-tailed t-test with Welch’s Correction; **Figure 3E**). Despite this large deletion of TH, FAA developed normally and did not differ from controls at any time point (p > 0.05; mixed effects analysis with Sidak’s multiple comparisons test; **Figure 3F**). Total activity was reduced on days 14 and 21 of the study (Day –7, p = 0.999; Day 0, p = 0.965; Day 7, p = 0.891; Day 14, p = 0.0412; Day 21, p = 0.0407; Day 28, p = 0.804; mixed effects analysis with Sidak’s multiple comparisons test; **Figure S3G**), but raw premeal activity remained similar between groups (p > 0.05; unpaired two-tailed t-test with Welch’s Correction; **Figure S3H**). Foxp2-TH-cKO mice were smaller than littermate controls throughout the study (**Figure S3I**), although normalized body weight during caloric restriction was largely comparable between genotypes (Day –7, p = 0.930; Day 0, p > 0.999; Day 7, p = 0.709; Day 14, p = 0.242; Day 21, p = 0.018; Day 28, p = 0.095; mixed effects analysis with Sidak’s multiple comparisons test; **Figure S3J**). Food intake was similar between groups (**Figure S3K**), although body weight–normalized intake again showed a mild increase in cKO mice (p = 0.0304; unpaired two-tailed t-test with Welch’s Correction; **Figure S3L**).

*Ntsr1^Cre^;Th^Flox^* cKO (Ntsr1-TH-cKO) mice showed 41 ± 15 TH+ SN neurons, significantly fewer than the 172 ± 10 observed in controls (p < 0.0001; unpaired two-tailed t-test with Welch’s Correction; **Figure 3G-H**). Despite this large deletion of TH, FAA developed normally and remained similar to controls at most time points, with a difference detected only on day 14 (Day –7, p = 0.973; Day 0, p = 0.920; Day 7, p > 0.999; Day 14, p = 0.043; Day 21, p = 0.183; Day 28, p = 0.650; mixed effects analysis with Sidak’s multiple comparisons test; **Figure 3I**). Total locomotor activity was reduced throughout the study (p _≤_ 0.005; mixed effects analysis with Sidak’s multiple comparisons test; **Figure S3M**). Raw premeal activity differed between groups at later time points (Day –7, p > 0.999; Day 0, p > 0.999; Day 7, p = 0.263; Day 14, p = 0.0001; Day 21, p < 0.0001; Day 28, p < 0.0001; mixed effects analysis with Sidak’s multiple comparisons test; **Figure S3N**). Body weights were largely similar between Ntsr1-TH-cKO and controls, except on day 0 (Day –7, p = 0.0849; Day 0, p = 0.0393; Day 7, p = 0.180; Day 14, p = 0.360; Day 21, p = 0.395; Day 28, p = 0.374; mixed effects analysis with Sidak’s multiple comparisons test; **Figure S3O**), and normalized body weights showed comparable weight loss during caloric restriction (**Figure S3P**). Food intake did not differ between genotypes (p = 0.7566; unpaired two-tailed t-test with Welch’s Correction; **Figure S3Q**), and this remained true after normalization to body weight (p = 0.541; unpaired two-tailed t-test with Welch’s Correction; **Figure S3R**).

*Sox6^Cre^;Th^Flox^* cKO (Sox6-TH-cKO) mice also exhibited a strong reduction in TH^+^ neurons (45 ± 4 vs. 171 ± 10; p = 0.0024; unpaired two-tailed t-test with Welch’s Correction; **Figure 3J-K**). Despite this deletion, FAA developed normally and was comparable to controls across the study (p > 0.05; mixed effects analysis with Sidak’s multiple comparisons test; **Figure 3L**). Total activity was reduced only on the final day of the experiment (Day –7, p = 0.588; Day 0, p = 0.198; Day 7, p = 0.157; Day 14, p = 0.098; Day 21, p = 0.206; Day 28, p = 0.0189; mixed effects analysis with Sidak’s multiple comparisons test; **Figure S3S**). Raw premeal activity differed between groups on days 21 and 28 (Day –7, p > 0.999; Day 0, p = 0.999; Day 7, p = 0.594; Day 14, p = 0.177; Day 21, p = 0.025; Day 28, p = 0.0019; mixed effects analysis with Sidak’s multiple comparisons test; **Figure S3T**). Sox6-TH-cKO mice weighed significantly less than controls throughout the study (Day –7, p = 0.0001; Day 0, p = 0.0004; Day 7, p = 0.0011; Day 14, p = 0.0069; Day 21, p = 0.0184; Day 28, p = 0.0239; mixed effects analysis with Sidak’s multiple comparisons test; **Figure S3U**). However, normalized body weights during caloric restriction were similar between groups (**Figure S3V**). While food intake differed between groups (p = 0.0012; unpaired two-tailed t-test with Welch’s Correction; **Figure S3W**), it remained comparable when normalized to body weight (p = 0.9456; unpaired two-tailed t-test with Welch’s Correction; **Figure S3X**).

Finally, *Slc17a6^Cre^;Th^Flox^* cKO (Vglut2-TH-cKO) mice showed a similarly strong reduction in TH^+^ neurons (58 ± 23; p < 0.0001; unpaired two-tailed t-test with Welch’s; **Fig. 3M–N**) due to the developmental expression and recombination of *Slc17a6^Cre^*.^44^ Despite this deletion, FAA developed normally in Vglut2-TH-cKO mice and was comparable to controls at most time points, with a difference detected on day 14 (Day –7, p > 0.999; Day 0, p = 0.995; Day 7, p = 0.948; Day 14, p = 0.0037; Day 21, p = 0.0559; Day 28, p = 0.436; mixed effects analysis with Sidak’s multiple comparisons test; **Figure 3O**). Total locomotor activity was reduced throughout the study (Day –7, p = 0.0089; Day 0, p = 0.001; Day 7, p = 0.0005; Day 14, p < 0.0001; Day 21, p < 0.0001; Day 28, p < 0.0001; mixed effects analysis with Sidak’s multiple comparisons test**; Figure S3Y**). Raw premeal activity differed between groups from day 7 onward (Day –7, p = 0.999; Day 0, p > 0.999; Day 7, p = 0.041; Day 14, p < 0.0001; Day 21, p < 0.0001; Day 28, p < 0.0001; mixed effects analysis with Sidak’s multiple comparisons test; **Figure S3Z**). Body weight was similar between groups across the study (**Figure S3AA**), although normalized body weight differed at later time points (Day –7, p = 0.996; Day 0, p > 0.999; Day 7, p = 0.437; Day 14, p = 0.0057; Day 21, p = 0.0004; Day 28, p < 0.0001; mixed effects analysis with Sidak’s multiple comparisons test; **Figure S3AB**). Food intake was similar between genotypes (p = 0.4131; unpaired two-tailed t-test with Welch’s Correction; **Figure S3AC**) and remained similar when normalized to body weight (p = 0.2644; unpaired two-tailed t-test with Welch’s Correction**; Figure S3AD**).

Across all five conditional knockout models, overall locomotor activity was reduced to varying degrees, with the strongest reductions observed in *Ntsr1^Cre^;Th^Flox^* and *Slc17a6^Cre^;Th^Flox^*cKO mice (**Figure S3M, Y**), whereas *Crhr1*, *Foxp2*, and *Sox6^Cre^;Th^Flox^*cKO mice showed more modest changes (**Fig. S3A, G, S**). Differences in raw premeal activity were most evident in the *Ntsr1-, Sox6*, and *Slc17a6^Cre^;Th^Flox^*cKO lines (**Figure S3N, T, Z**), whereas *Crhr1 and Foxp2^Cre^;Th^Flox^*cKO mice tracked closely with controls across the 28-day caloric restriction protocol (**Figure S3B, H**). Despite substantial reductions in TH expression across each population, all five lines retained robust FAA, indicating that anticipatory locomotion does not depend on the majority of molecularly defined nigrostriatal dopamine neuron populations. Body weight and food intake were also broadly similar between genotypes, although several lines displayed lower absolute body weight and mild hyperphagia when intake was normalized to body weight (**Figure S3**).

### Conditional Deletion of Tyrosine Hydroxylase in Calb1-Cre Dopamine Neurons Severely Impairs FAA

The persistence of FAA across five independent cKO lines was unexpected and suggested that anticipatory locomotion could depend on a comparatively small nigrostriatal dopamine neuron population. We next employed the *Calb1^Cre^* driver,^45^ which targets multiple DAT⁺ neuronal subtypes in the classification scheme of Poulin et al. (2014).^29^ In *Calb1^Cre^;Th^Flox^* cKO (Calb1-TH-cKO) mice, histological analysis revealed only a modest reduction in TH^+^ neurons within the SN, with a 23.2% decrease relative to controls (p = 0.0072, unpaired two-tailed t-test with Welch’s Correction; **Figure 4A-B**). Body weights were normalized to each animal’s starting weight on day 0 of CR, and both control and cKO groups lost comparable fractions of their initial weight throughout the CR period, with no significant differences at any time point (p > 0.05, mixed effects analysis with Sidak’s multiple comparisons test; **Figure 4C**). Consistent with this, average body weights did not differ between groups during the CR protocol (p > 0.05; mixed effects analysis with Sidak’s multiple comparisons test; **Figure S4A**). Food intake was also unaffected by the knockout (**Figure S4B**). Control animals consumed 0.17 ± 0.008 g/g, whereas cKO mice consumed 0.166 ± 0.009 g/g of body weight (p = 0.7434, unpaired two-tailed t-test with Welch’s Correction; **Figure 4D**).

**Figure 4.**
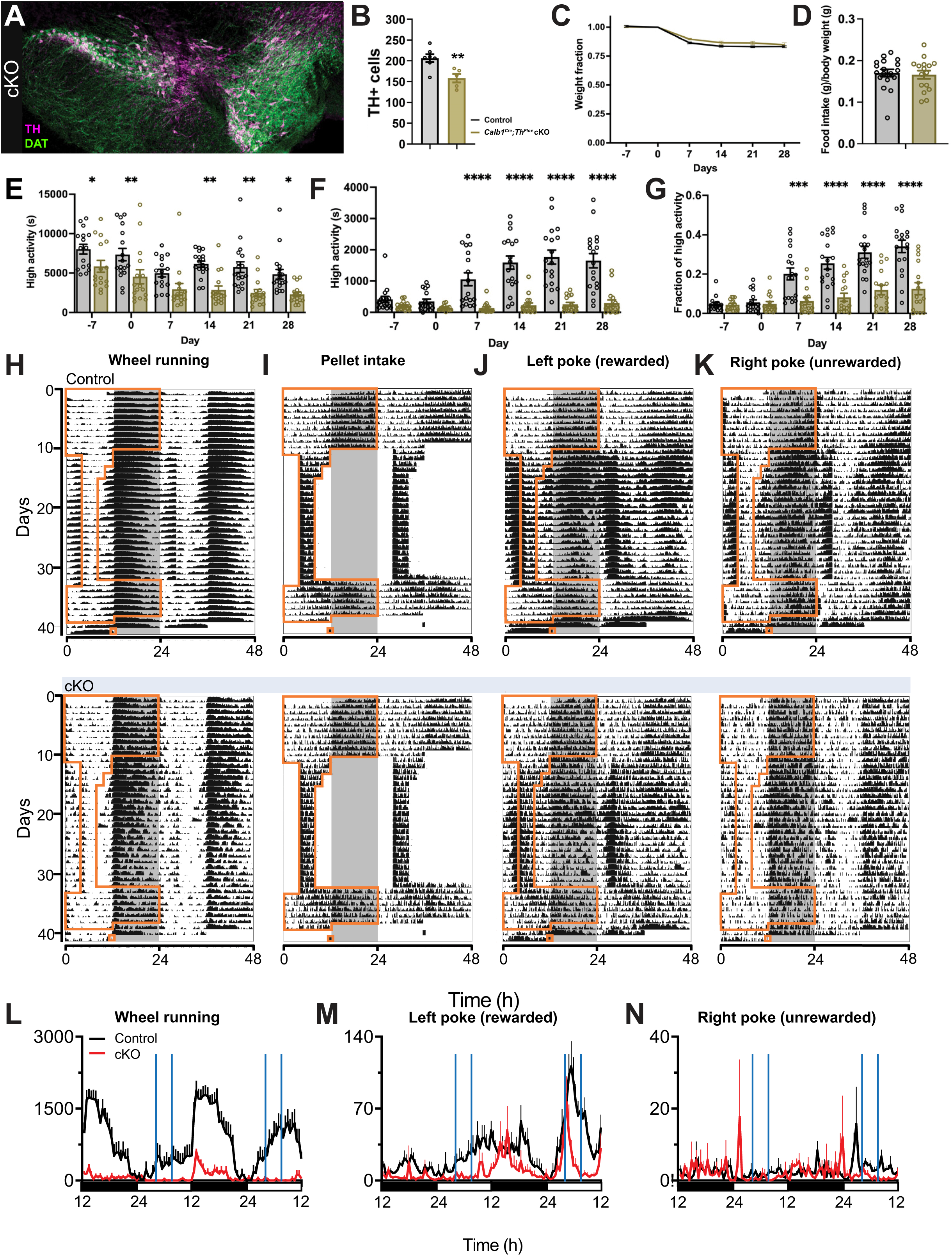
Conditional Deletion of Tyrosine Hydroxylase in *Calb1-Cre* Dopamine Neurons Severely Impairs FAA and AAV-TH Injection into SN Fails to Rescue FAA. A) Confocal imaging of immunofluorescence antibody staining of Th (magenta) and DAT (green) in the midbrain of *Calb1^Cre^;Th^Flox^* cKO midbrain tissue. B) Quantification of TH staining in the SN of WT (grey) and *Calb1^Cre^;Th^Flox^* cKO (gold) midbrain tissue. C) Normalized body weight of WT and *Calb1^Cre^;Th^Flox^* cKO mice over the course of the study; weight was normalized by dividing the weekly weight by the Day 0 weight, which was taken prior to beginning CR. The weight fraction was similar between the two groups at every time point. D) Normalized food intake of WT and *Calb1^Cre^;Th^Flox^* cKO mice. Food intake was normalized by dividing average 24-hour AL intake by the animal’s body weight. E) Total daily high activity in seconds of WT (grey) and *Calb1^Cre^;Th^Flox^* cKO (gold) mice throughout the course of the study. F) High activity in seconds during the 3-hour premeal window throughout the course of the study. G) Normalized high activity during the 3-hour premeal window; data is normalized by dividing the seconds of high activity in the premeal window by the total daily high activity. H-K) Group-averaged double-plotted ethograms for WT and *Calb1^Cre^;Th^Flox^* cKO mice over a 40-day period for wheel running activity (H), pellet intake (I), rewarded left nose pokes (J), and unrewarded right nose pokes are shown. Area within brown borders indicates periods of food availability, and dark phase is indicated by gray background. L-M) Group average profiles for wheel running activity, rewarded left nose pokes, and unrewarded right nose pokes during 48 h food deprivation. Blue vertical lines indicate onset and offset of food availability during previously restricted feeding schedule. Light-dark cycle indicated as black and white at the bottom of each graph.

Under AL feeding conditions, Calb1-TH-cKO mice exhibited normal daily rhythmicity, with minimal daytime activity like controls (**Figure S4C**). However, total locomotor activity was significantly lower in cKO mice than in controls at most time points (Day -7, p = 0.0424; Day 0, p = 0.0082; Day 7, p = 0.1277; Day 14, p = 0.0013; Day 21, p = 0.0014; Day 28, p = 0.0228; mixed effects analysis with Sidak’s multiple comparisons test; **Figure 4E**). Despite maintaining normal daily activity patterns under AL feeding, Calb1-TH-cKO mice displayed a profound impairment in FAA during scheduled feeding. Prior to CR, both groups exhibited similarly low levels of premeal activity, with no significant difference between controls (412 ± 96 s) and cKO mice (day –7, 212 ± 34 s; p = 0.8336; mixed effects analysis with Sidak’s multiple comparisons test; **Figure 4F**). By day 7 of CR, control mice showed a robust increase in premeal activity (1057 ± 206 s), whereas cKO mice remained largely inactive (132 ± 44 s; p < 0.0001; mixed effects analysis with Sidak’s multiple comparisons test). This divergence persisted and widened through day 28, when control mice reached 1651 ± 226 s of premeal activity compared to 295 ± 102 s in cKO mice (p < 0.0001; mixed effects analysis with Sidak’s multiple comparisons test; **Figure 4F**). Because Calb1-TH-cKO mice exhibited reduced overall locomotor activity, we next examined normalized FAA, calculated as the fraction of total daily activity occurring in the premeal window. This analysis confirmed that FAA remained significantly reduced in cKO mice (Day -7, p > 0.999; Day 0, p > 0.999; Day 7, p = 0.0003; Day 14, p < 0.0001; Day 21, p < 0.0001; Day 28, p < 0.0001; mixed effects analysis with Sidak’s multiple comparisons test; **Figure 4G**). Thus, the deficit in FAA cannot be explained solely by reduced overall locomotor activity.

To determine whether the absence of FAA reflected a generalized failure to arouse from sleep, we delivered a mild arousal stimulus—removal of the cage lid accompanied by gentle cage movement—two hours before the scheduled feeding time. During AL feeding, both control and Calb1-TH-cKO mice showed minimal responses to this stimulus (WT: 278.0 ± 42.6 s; cKO: 118.3 ± 37.2 s; p = 0.0183; unpaired two-tailed t-test with Welch’s Correction; **Figure S4D, S4F**). However, when tested again after 23 days of timed CR, control mice displayed robust anticipatory activity even in the absence of stimulation, whereas cKO mice remained largely inactive (**Figure S4E**). When the anticipatory cue was delivered on day 23, control mice exhibited sustained activity throughout the two-hour post-cue period (2,780.0 ± 533.6 s), representing approximately a tenfold increase over their baseline response during AL feeding **(Figure S4F-G).** In contrast, Calb1-TH-cKO mice displayed only 285.0 ± 80.99 s of activity, the majority of which (75.9 ± 8.2%) occurred within the first 25 minutes after the cue, after which activity rapidly declined **(Figure S4F-G).**

Because *Calb1^Cre^* is not restricted to midbrain dopamine neurons, we examined additional catecholaminergic populations. Calb1-TH-cKO exhibited significant reductions in TH^+^ neurons within the hypothalamic A11 and A12 dopamine populations, whereas TH expression in A13 neurons was unchanged (**Figure S5A-B**). In contrast, TH+ neuron numbers in the locus coeruleus were unaffected (**Figure S5A-B**), indicating that *Calb1^Cre^* does not substantially target noradrenergic neurons. To assess adult *Calb1^Cre^* activity within hypothalamic dopamine populations, we injected *Calb1^Cre^*;Ai6 mice with AAV9-PHP.eB-CAG-FLEX-tdTomato. Despite robust lineage lab**e**ling in Ai6 mice, little tdTomato expression was observed in A11 or A12 regions (**Figure S5C**), suggesting limited adult Cre activity in these populations.

### AAV-TH Injection into SN Fails to Rescue FAA in Calb1^Cre^;Th^Flox^ cKO Mice

Given the marked suppression of FAA in Calb1-TH-cKO mice, we attempted a targeted rescue by restoring TH expression in the SN, as previously performed in DAT-TH-cKO mice (**Figure 2**). To this end, we delivered stereotaxic intracranial (IC) injections of either AAV8-DIO-TH-RFP (n = 10) or control AAV9-DIO-GFP (n = 9) into the SN of Calb1-TH-cKO mice. FAA was not restored in AAV-DIO-TH-RFP injected mice, nor was it in negative controls, as expected. Both groups exhibited similarly low total daily locomotor activity throughout the 28-day CR protocol, with no significant differences between viral groups at any time point (p > 0.05; mixed effects analysis with Sidak’s multiple comparisons test; **Figure S6J**). Consistent with this, neither time (p = 0.561) nor viral genotype (p = 0.738) significantly affected total activity. FAA similarly failed to recover following viral TH restoration. Premeal activity remained statistically indistinguishable between groups across all days (Day –7: p = 0.998; Day 0: p = 0.814; Day 7: p = 0.999; Day 14: p = 0.533; Day 21: p = 0.827; Day 28: P = 0.999; mixed effects analysis with Sidak’s multiple comparisons test; **Figure S6K)**. The same pattern was observed for normalized premeal activity (Day –7: p = 0.909; Day 0: p = 0.363; Day 7:p > 0.999; Day 14: p > 0.999; Day 21: p = 0.716; Day 28: p = 0.778; mixed effects analysis with Sidak’s multiple comparisons test; **Figure S6L**), confirming a lack of behavioral rescue.

To determine whether viral delivery successfully restored TH expression, we next examined midbrain sections using immunohistochemistry. RFP+/TH+ double-positive neurons were sparse in the SN of AAV-DIO-TH-RFP injected mice (**Figure S6A-B**). Across five SN sections, the total number of RFP+ cells averaged 11.5 ± 2.8 per section, and we observed an average of 4.1 ± 1.1 RFP+/TH+ neurons per section, corresponding to approximately 20.5 double-positive neurons per brain **(Figure S6D).** This number was far below the ∼50 rescued neurons previously observed in *Slc6a3^Cre^;Th^Flox^*cKO AAV rescue experiments. Notably, approximately 64.1% of RFP+ cells lacked TH expression, suggesting either non-dopaminergic viral transduction or incomplete TH re-expression. The number of GFP+ cells observed in control injections averaged to 8.7 ± 2.3 cells per section (**Figure S6C**). Importantly, the total number of TH+ neurons in the SN was nearly identical between groups (114.1 ± 8.4 vs. 110.5 ± 6.7 cells per section; P = 0.748, unpaired two-tailed t-test with Welch’s Correction; **Figure S6E**), indicating that viral delivery did not significantly restore TH expression.

We also monitored body weight and food intake following viral delivery. Raw body weights did not differ between groups across the CR protocol (Day –7: p = 0.994; Day 0: p = 0.989; Day 7: p = 0.984; Day 14: p = 0.909; Day 21: p = 0.698; Day 28: p = 0.829; mixed effects analysis with Sidak’s multiple comparisons test; **Figure S6F**), and normalized body weights were similarly indistinguishable (Day –7: p > 0.999; Day 7: p = 0.9991; Day 14: p = 0.946; Day 21: p = 0.857; Day 28: p = 0.824; mixed effects analysis with Sidak’s multiple comparisons test; **Figure S6G**). Food intake was modestly higher in the AAV-DIO-TH-RFP group (4.0 ± 0.199 g) compared with controls (3.4 ± 0.172 g; p = 0.0397, unpaired two-tailed t-test with Welch’s correction; **Figure S6H**). However, this difference disappeared when intake was normalized to body weight (0.162 ± 0.009 g/g vs. 0.144 ± 0.010 g/g; p = 0.215; unpaired two-tailed t-test with Welch’s Correction; **Figure S6I**).

Due to the failure of AAV-mediated TH rescue to restore FAA, together with minimal RFP and GFP reporter expression in the SN, we characterized the distribution of *Calb1* mRNA and Calbindin1 protein across the ventral midbrain **(Figure S7).** Immunofluorescence labeling revealed limited Calbindin1 expression across SN and VTA (**Figure S7A**). In the SN, TH⁺ neurons exhibited minimal Calbindin1 co-expression, as only 16.8% of dopaminergic neurons were double positive for both markers (18.2 ± 2.2 TH+/Calb1+ cells vs. 90.1 ± 5.7 TH+ cells per section; **Figure S7B)**. In contrast, Calbindin1 expression was more prominent within the VTA, where there was robust co-labeling in 57.3% of dopaminergic neurons (76.9 ± 7.5 TH+/Calb1+ cells vs. 57.3 ± 6.5 TH+ cells per section; **Figure S7B**). We next validated these findings using fluorescence *in situ* hybridization for *Th* and *Calb1* transcripts (**Figure S7C**), which recapitulated the protein-level results. *Calb1* mRNA was sparse within TH⁺ neurons in the SN but was readily detected in a subset of VTA dopaminergic neurons, indicating that the observed distribution is not attributable to antibody sensitivity. To confirm functional *Calb1^Cre^* recombinase activity in the VTA, we injected a Cre-dependent “tet bow”^46^ AAV8 cocktail (CAG-DIO-tTA, TRE-DIO-mNeonGreen, TRE-DIO-mTurquoise2, and TRE-DIO-mRuby2). Fluorescent labeling was observed in the VTA, consistent with active Cre recombination in adulthood (**Figure S7D**). Surprisingly, analysis of a genetic cross between *Calb1^Cre^* to Cre-dependent reporter line (Ai9) showed no reporter activity in TH⁺ neurons in SN or VTA (**Figure S7E**). To further distinguish developmental lineage from adult Cre activity, we systemically delivered AAV9-PHP.eB-CAG-FLEX-tdTomato to adult *Calb1^Cre^*;Ai6 mice. Whereas Ai6 reporter expression was prominent in the VTA and identified sparse developmental recombination in the SN, adult FLEX reporter expression was nearly absent from SN dopamine neurons and remained largely restricted to the VTA (**Figure S7F-G**). Together, these findings further support the conclusion that *Calb1^Cre^* provides limited genetic access to the adult SN dopamine population targeted by the developmental knockout.

### Calb1^Cre^;Th^Flox^ cKO Mice Exhibit Impaired Anticipatory Locomotion but Preserved Food-Seeking Behavior

To determine whether the loss of FAA in Calb1-TH-cKO mice reflected impaired food-entrained timekeeping or a selective deficit in locomotor output, we evaluated anticipatory food-seeking behavior using a multi-modal behavioral paradigm combining voluntary wheel running, pellet intake, and operant nose-poking.^5,47^ Mice were monitored under AL feeding and daytime-restricted feeding conditions using the FED3 system.^48^

Under AL feeding conditions, both control and Calb1-TH-cKO mice exhibited strong nocturnal behavioral rhythms across all measured outputs, including wheel running, pellet intake, and nose-poking (**Figure 4H–K**). However, total wheel-running activity was markedly reduced in Calb1-TH-cKO mice compared with controls (**Figure S8A-C**). Despite this reduction in locomotor output, pellet intake and rewarded left nose-poking during the dark phase remained largely similar between genotypes, indicating preserved feeding behavior and operant performance **(Figure S8D-I).** During daytime-restricted feeding, control mice developed robust FAA characterized by increased wheel running, elevated rewarded nose-poking, and increased pellet intake in the hours preceding scheduled food availability (**Figure 4H-K**). In contrast, Calb1-TH-cKO mice failed to exhibit anticipatory wheel-running activity, and overall locomotor output remained low throughout the restricted feeding phase (**Figure 4H, L**). Despite the profound suppression of anticipatory locomotion, Calb1-TH-cKO mice retained significant anticipatory food-seeking behavior, as measured by rewarded nose-poking (**Figure 4J, M**). Although anticipatory nose-poking emerged more slowly and reached lower peak levels in cKO mice compared with controls, its persistence indicates that food-seeking behavior was less affected than anticipatory locomotion.

Consistent with this interpretation, anticipatory nose-poking disappeared rapidly when animals were returned to AL feeding, and diurnal pellet intake gradually shifted back to the nighttime active phase (**Figure 4I-J**). Furthermore, during a subsequent 48-hour food deprivation, anticipatory nose-poking reappeared at approximately the same circadian time as the previously scheduled feeding window in both control and cKO mice (**Figure 4M–N**). In contrast, anticipatory wheel-running activity did not re-emerge in cKO mice. Unrewarded right nose-poking also showed anticipatory increases in both groups, although the magnitude of this response was lower in cKO mice (**Figure 4K, N**). Together, these findings demonstrate that *Calb1*-expressing dopamine neurons are required for the locomotor expression of FAA but are not necessary for the circadian prediction of food availability.

## Discussion

The neural mechanisms that enable animals to anticipate daily food availability independently of the light-entrained circadian clock have remained elusive despite decades of investigation. Although dopamine signaling has long been implicated in food anticipatory activity (FAA), whether dopamine neurons participate directly in food-entrained timing or instead mediate its behavioral expression has remained unresolved. Using systematic genetic dissection of molecularly defined dopamine neuron populations, we demonstrate that FAA does not require most major nigrostriatal dopamine neuron populations. Instead, a relatively small Calbindin1-lineage dopamine population is selectively required for anticipatory locomotion. Remarkably, loss of dopamine synthesis in this population profoundly suppresses anticipatory locomotor activity while sparing substantial anticipatory food-seeking behavior. These findings identify a previously unrecognized dopaminergic population involved in food anticipation and suggest that distinct behavioral components of anticipation can be differentially regulated.

Conditional deletion of *Th* in DAT-expressing neurons confirmed a critical role for dopamine synthesis in FAA. *Slc6a3^Cre^;Th^Flox^*cKO mice failed to develop FAA under scheduled feeding, consistent with earlier pharmacological and genetic studies implicating dopamine signaling in this behavior.^15,16,27^ Importantly, restoring TH expression in a limited number of SN dopamine neurons was sufficient to rescue FAA despite persistent hypoactivity and unchanged food intake. These findings demonstrate that dopamine production in a surprisingly small subset of midbrain neurons is sufficient to support anticipatory locomotion, consistent with our prior results in *Pitx3^Cre^* mutants,^28^ but they do not establish whether dopamine neurons themselves function as FEOs or instead act downstream of circadian prediction.^49^ Our systematic Cre- based deletion screen provides strong evidence for the latter interpretation. Deletion of *Th* using five independent *Cre* driver lines—*Crhr1, Foxp2, Ntsr1, Sox6, and Slc17a6*—each eliminated large fractions of SN dopamine neurons, in some cases exceeding 75–80% of the population. Despite this extensive loss, FAA developed and persisted in all lines. These findings indicate that neither the bulk of nigrostriatal dopamine neurons nor several major molecularly defined dopamine subtypes are required for FAA. Thus, robust anticipatory locomotion can emerge despite extensive loss of several major dopaminergic populations, in contrast to what may have been expected based on other studies implicating dopamine signaling in FAA.^3,16,24–27,49^

A key caveat in interpreting these deletion experiments is the possibility of incomplete recombination if Cre expression is transient or weak during development or thereafter. For example, dual-recombinase studies indicate that ∼98% of SN dopamine neurons express *Slc17a6^Cre^* during development,^44^ whereas our *Slc17a6^Cre^*–driven *Th* deletion reduced TH expression by only ∼70% in the SN. By contrast, *Crhr1^Cre^*produced a robust ∼82% loss of TH. Remarkably, even these mice—despite being underweight and requiring food on the cage floor—exhibited strong FAA (a similar phenotype to *Slc6a3^Cre^*-mediated cKO). These findings echo studies of *Pitx3^Cre^* hypomorphic mice, which have severely depleted SN dopamine neurons yet retain FAA and metabolic adaptation to scheduled feeding.^28^ Together, these observations suggest that *Pitx3*-negative and *Crhr1*-, *Foxp2*-, *Ntsr1-, Sox6-*, and *Slc17a6^Cre^*–expressing SN dopamine populations are not required for FAA, supporting the idea that a smaller and more specific dopamine subpopulation mediates this behavior.^28^

In striking contrast to these preserved phenotypes, deletion of *Th* using *Calb1^Cre^* produced a profound and persistent loss of FAA despite affecting only a modest fraction of SN dopamine neurons. *Calb1^Cre^;Th^Flox^*cKO mice maintained normal feeding behavior, body-weight regulation, and nocturnal activity patterns under AL conditions, indicating that the FAA deficit cannot be explained by metabolic impairment, generalized hypoactivity, or circadian disruption. Instead, these mice failed specifically to increase locomotor activity prior to scheduled feeding and were unable to sustain anticipatory activation even in response to external arousal cues. This disproportionate behavioral effect identifies Calbindin1-lineage dopamine neurons as a critical bottleneck linking food-entrained signals to anticipatory locomotion. Importantly, the absence of FAA in Calb1-TH-cKO mice did not reflect a failure of food-entrained timekeeping. These mice displayed robust anticipatory food-seeking behavior, as measured by operant nose-poking for food, during restricted feeding. Anticipatory nose-poking disappeared upon return to AL feeding and re-emerged during food deprivation at the previously scheduled mealtime, demonstrating intact circadian prediction of food availability. Thus, predictive information about mealtime is preserved, but its translation into locomotor output is selectively disrupted. These findings provide genetic evidence that distinct behavioral components of food anticipation can be dissociated.

Attempts to rescue FAA in *Calb1^Cre^;Th^Flox^*cKO mice via adult AAV-mediated restoration of TH expression in the SN were unsuccessful, in contrast to the robust behavioral rescue observed in *Slc6a3^Cre^;Th^Flox^*cKO mice. This failure is unlikely to reflect insufficient overall TH levels, as total TH^+^ neuron numbers in the *Calb1^Cre^;Th^Flox^*cKO mice were comparable to or greater than those in successfully rescued *Slc6a3^Cre^;Th^Flox^*cKO mice. Instead, these findings suggest that restoring TH in the correct dopaminergic subpopulation, rather than overall TH levels, is critical for rescuing FAA. One likely explanation is that Calb1Cre does not efficiently target the relevant SN dopamine neurons in adulthood. A related caveat is that *Calb1^Cre^*-mediated recombination is not restricted to the SN and also targets subsets of hypothalamic dopamine neurons (**Figure S5A-B**). However, *Calb1^Cre^*does not significantly target A13 dopamine neurons or locus coeruleus norepinephrine neurons, and adult Cre activity within A11/A12 populations is minimal as assessed by systemic Cre-dependent reporter expression (**Figure S5C**). Consistent with this, *Calb1^Cre^* expression appears sparse in the adult SN, limiting its utility for adult-specific manipulations. Evidence for this includes: (1) sparse SN labeling in adult brains by immunohistochemistry (while VTA labeling remains robust); (2) weak *Calb1 in situ* hybridization signal in the SN; (3) minimal Cre-dependent AAV reporter expression (AAV-DIO-GFP) in the SN of adult mice; and (4) limited TH restoration following AAV-DIO-TH-RFP injection, largely restricted to the *pars lateralis*. Together, these observations indicate that the failure to rescue FAA reflects incomplete targeting of a functionally critical dopaminergic subpopulation, rather than insufficient TH expression per se.

Taken together, our findings are more consistent with a distributed model of food-entrained timing than with a unitary dopamine-based FEO. Under this framework, predictive signals generated by one or more food-entrainable oscillators converge on a specialized dopaminergic population that is required for the expression of anticipatory locomotion. Rather than serving as the primary timing mechanism itself, this population may function as a critical output node linking food-entrained prediction to behavioral activation. Notably, Calbindin1-positive SN dopamine neurons preferentially innervate the dorsomedial striatum,^50^ a region implicated in internally guided action selection and motivational state-dependent behavior.^51,52^ Furthermore, it was demonstrated that a specific medial pre-frontal cortical input to the DS can promote running behavior specifically during conditions of food restriction, suggesting that there may be multiple inputs to the DS that regulate pre-meal activity levels.^53^ Our working hypothesis is that anticipatory locomotion may be mediated through a restricted striatal circuit that couples circadian prediction to motor activation. These findings raise the possibility that multiple DMS-targeting circuits converge to regulate pre-meal arousal. Future studies mapping the inputs to and outputs of Calbindin1-positive SN dopamine neurons will be essential for identifying the upstream circuitry responsible for food-entrained timing signals.

## Methods

### Ethics Statement

The experiments described herein were approved by the California State Polytechnic, Pomona Institutional Animal Care and Use Committee (IACUC) under protocols: 13.025, 13.029, 16.029, 17.003, 20.013, and 20.014, or IACUC at UT Southwestern Medical Center under protocol: 2016 101376-G.

### Animal Husbandry

Mice used in experiments were maintained in a 12:12 light:dark cycle throughout all portions of study. Zeitgeber Time (ZT) 0 was defined as lights on, whilst ZT 12 was lights off, by convention. Mice were housed in static microisolator cages using layered Sani-Chip bedding (Envigo, 7090), plastic or paper shelter structures, and cotton or paper nestlets. Shelter and nestlets were excluded only when being video recorded to measure behavior. Cages included a wire rack to provide access to a water bottle and standard rodent chow (Envigo, 2018 Teklad 18% Protein Rodent Diet) with caloric density of 3.1 kcal/g. Ambient temperatures were maintained between 22-24°C with humidity between 20-45%. To promote the survival of *Slc6a3*Cre;ThFlox cKO mice, crushed chow along with Ensure® (Abbot Laboratories) or Nutrical® (Vetoquinol, USA) was added to the bottom of the cage. This was replaced every other day until the DAT KO mice reached 6 weeks old. When mice were at least 10 weeks old they were single-housed for 5-7 days with AL access to food and water prior to being placed on CR.

### Breeding scheme for generating conditional knockout lines

*Th^Flox^* and *Th^null^* (Slc6a3^tm1(cre)Xz^/J, Stock #020080 and B6;129-*Dbh^tm2(Th)Rpa^ Th^tm1Rpa^/J*, #009688) mouse lines were provided by Dr. Richard Palmiter (Univ. of Washington, Seattle). The *Th^null^* allele contains a localized deletion of *Th,* which is available from the Jackson Laboratory (Bar Harbor, ME; stock #009688). The *Th^Flox^*allele has loxP sites flanking the first exon of *Th*.^33^ *Slc6a3^Cre^*-driver mice were obtained from the Jackson Laboratory (Bar Harbor, ME; stock #020080); this line contains a gene insertion of *Cre* recombinase upstream of *Slc6a3*.^35^ *Calb1^Cre^*-driver mice were obtained from The Jackson Laboratory (Bar Harbor, ME; stock #028532); this allele contains an internal ribosomal entry site (IRES).^45^ *Foxp2^Cre^*-driver mice were obtained from The Jackson Laboratory (Bar Harbor, ME; stock #030541); this allele contains an IRES. *Crhr1^Cre^*-driver mice were kindly supplied to us by Dr. Larry Zweifel (University of Washington, Seattle) and contains an IRES-Cre:GFP allele.^54^ *Slc17a6^IRES-Cre^* knock-in mice were obtained from Jackson Laboratory (stock# 016963)^43^ and was also an IRES allele. *Ntsr1^Cre^* mice were derived from the breeding colony of Dr. Gina Leinninger (Michigan State University) and are also available from Jackson laboratory (stock #033365); this is also an IRES allele.^41^ *Sox6^Cre^* mice were obtained from the laboratory of Rajeshwar Awatramani of Northwestern University, Feinberg School of Medicine (Supp. Table I).^42^ Progeny that was Cre+ and either *Th*^+/-^ or *Th*^+/Flox^ were backcrossed to the parental line containing one loxP flanked (“Flox”) allele to generate cKO mice. For genotyping, tail clips were sent to Transnetyx (Cordova, TN) for qPCR analysis. Genetic purity analysis for all strains used were >97% C57BL/6 by Transnetyx (Cordova, TN).

### Food Intake, Caloric Restriction, and Home-Cage Behavioral Measurements

Mice were individually housed and measured for daily food intake, or the amount of food consumed by a mouse in a 24-hour period, while having free or *ad libitum (*AL) access to standard rodent chow. Daily food intake data were used to calculate the 60% CR amount each mouse would be given throughout the FAA study, which were liable to adjustment to maintain a body weight of around 85% of their total body weight at the beginning (day 0) of the study. Over the course of the study, body weights (in grams) were measured weekly. Food intake tests were conducted before starting mice on CR at ages of ∼9-10 weeks. To calculate food amounts that would be served as the 60% CR food values per mouse cohort, food intakes were measured per mouse via the placement of ∼40g of rodent chow in food bins and on floors of home cages, then measuring the remaining mass of chow 24h after. These values were collected over ∼48-72 hours and were then averaged per group and 60% of the daily average was determined as the daily CR value. Mice were then fed 60% of their daily caloric intake daily for 28 days at ZT 6 starting on Day 0, the start of CR. After Day 28 of study, mice were euthanized by CO_2_ narcosis, and were transcardially perfused with phosphate buffer followed by 4% paraformaldehyde.

### Home-cage behavioral measurements

Home-cage behaviors were measured and recorded by video recording mice from a perpendicular angle to their home cages. Videos were then analyzed using an automated behavior recognition software, HomeCageScan 3.0,^37,54^ which annotates the following behaviors: remain low, pause, twitch, awaken, distance traveled, turn, sniff, groom, food bin entry, chew, drink, stretch, unassigned behaviors, hanging, jumping, rearing, and walking. HomeCageScan outputs its analyzed data into an excel sheet format of twenty-four one-hour bins to quantify the temporal structure of measured activities. The sum high activity only accounts for hanging, jumping, rearing, and walking, as these behaviors are designated as high activity; all other scored behaviors were not used. Total high activity was then determined by summation of high activity bins, while FAA ratios were calculated by dividing the final 3h of activity over the total high activity of each mouse. For home-cage behavioral measurements, behaviors were recorded starting on Day -7 at ∼9-10 weeks of age. During filming periods, Sani-Chip bedding was minimized to 100mL per cage and all other enrichment was removed. Dim red LED lighting was provided with high power 42 SMT Red LED PAR38 from LEDwholesalers.com.

### Automated behavioral and feeding monitoring (UT Southwestern)

A subset of this study involving behavioral monitoring was conducted at the University of Texas Southwestern Medical Center (UTSW). Twenty male and female *Calb1^Cre^;Th^Flox^* mice and control littermates (C57BL/6J background) were sent to UTSW, where they were tested across three cohorts: Mice of Cohorts 1 (n=7; 4 males, 3 females) and Cohort 3 (n=7; 5 males, 2 females) were bred at California State Polytechnic University, Pomona, and were shipped to UTSW. Cohort 2 (n=6; 3 males, 3 females) was bred on-site at UTSW. See Table S1 for genotype, age and sex of each mice used. At experimental onset, mice were approximately 9 weeks (Cohort 1), 5 weeks (Cohort 2), and 15 weeks of age (Cohort 3). Due to FED3 malfunctions during 42 days of recording period, 1 mouse in cohort 1 and 2 mice in cohort 2 were excluded from all plots and analysis.

### Housing and FED3 Configuration (UTSW)

Mice were individually housed in cages (14.5 × 32.5 × 13.0 cm) equipped with 11-cm diameter running wheels and woodchip bedding (Sani-Chips, PJ Murphy Forest Products). Animals were maintained under a 12-h light/12-h dark cycle (250 lux) within light excluding, ventilated cabinets. Temperature, humidity, and light intensity were recorded every 5 minutes using Chamber Controller software (ver. 4.104, Actimetrics, Wilmette, IL, United States). Feeding activity was measured using a Feeding Experimental Device 3 (FED3). The FED3 comprises two nose-poke ports; the left reward port and the right inactive port. Upon nose poking, mice were rewarded with a 20 mg grain-based pellet (Bio-Serv #F0163) from the left port, or no reward from the right port. To eliminate external time cues, internal LEDs and buzzers were disabled, and any maintenance was conducted at randomized intervals. was mounted on the side of each cage, with holes cut to allow nose-poke access for pellet delivery.

### Time Restricted Feeding Protocol and Data Analysis (UTSW)

In the first 10 days of the procedure, mice were allowed AL access to FED3 delivered food. Mice were then transitioned to a time restricted feeding schedule following a 16-h overnight fast (starting at ZT 12 on day 11), to a 4-h daytime feeding schedule during ZT 4-8. This included 2 days of 8-h access and 2 days of 6-h access, followed by 17 consecutive days of 4-h access. Following restricted feeding phase, mice were returned to an AL feeding schedule for 7 days, followed by a 48-h total food deprivation period beginning at ZT 12.

Wheel-running activity was recorded in 1-min bins via the ClockLab acquisition system (ver. 3.604, Actimetrics). ClockLab (ver. 6.1.15) was used to generate double-plotted group-average and individual ethograms (10-min bins) using percentile plots (quantiles = 50). 24-h group profiles (30-min bins) were generated for the final 7 days of the AL phase (Days 4–10) and the final 7 days of the RF phase (Days 26–32).

FAA and anticipatory nose-poking were quantified as the total events occurring during the 4-h window preceding food availability (ZT0 to ZT4). One *Calb1^Cre^;Th^Flox^* mouse failed to acquire the nose-poking task during the AL phase and was excluded from pellet intake analysis but retained for all other behavioral metrics. Statistical significance between genotypes was determined using the Mann-Whitney test.

### Surgical procedures

Stereotaxic delivery of AAV serotype 8 was used to restore TH in a Cre-dependent manner in *Slc6a3^Cre^* and *Calb1^Cre^*TH-cKO mice. AAV-DIO-TH-RFP was injected to restore TH and AAV-DIO-GFP was injected as a negative control. Mice underwent surgery at 8-12 weeks of age. Mice undergoing surgery were first induced in a sealed induction chamber with an oxygen flow rate of 1.5 lpm O2 and 5% Isoflurane. Once fully anesthetized, mice were placed onto a stereotaxic frame (Kopf, Model 1900 Stereotaxic Alignment System) and stabilized. The scalp was sterilized with three washes of chlorhexidine followed by isopropanol, and an incision was made to expose the skull. Excess fascia was removed with sterile cotton swabs to allow for full exposure of the skull. Once exposed, the skull was leveled (Kopf, Model 1905 Stereotaxic Alignment Indicator), and the drill was centered on bregma, the intersection of the coronal and sagittal sutures of the skull. From this point, holes were drilled in the skull using coordinates to specifically target the SN (x = +/-1.5mm, y = -3.6mm) or VTA (x = +/-0.5mm, y = -3.7mm). A microinjector with an attached glass pipet loaded with virus was slowly lowered into the brain to the proper depth (-4.0 mm). After waiting 5 min, bilateral injections of 750 nL of virus were given at a delivery rate of 10 nL/second, followed by a 10-min waiting period following each injection to optimize uptake of virus into surrounding tissue. Following injection, the skull was sealed with bone wax (Surgical Specialties, #309) and 0.1 mL topical anesthetic (Hospira, Marcaine 0.5% 5 mg/mL, PAA113011) was applied to the skull and incision site prior to sealing the incision with VetBond (3M, No. 1469SB). After isoflurane was turned off, an additional 0.5mL sterile saline was injected subcutaneously. Mice were transferred into a sterile home-cage placed beneath a heating lamp until consciousness returns.

Animals were weighed before and after the procedure to account for weight gain due to supplementary fluids. Animals were monitored for a minimum of three days following surgery and were given 0.1mL 0.5mg/mL ketoprofen and Nutri-Cal (Vetoquinol) daily; an additional 0.5mL saline SQ could also be provided at researcher’s discretion, based on animal condition. The protocol states that the humane endpoint following the procedure is 20% loss of body weight within the monitoring period, after which an animal would be euthanized.

*Th* cKO mice were selected for injection once aged 8-12 weeks, with a minimum weight of 16 g being required for an animal to be considered for surgery. Two separate Adeno-Associated Viruses (AAVs) that are encoded in a *Cre*-dependent manner created by Vector Biolabs (Malvern, PA) were utilized. The first construct (AAV DIO-TH-RFP) encodes a red fluorescent protein (RFP) conjugated with *Th* via P2A, a self-cleaving peptide that mediates ribosomal skipping during translation, to allow for consistent co-expression. The second construct (AAV DIO-GFP) encodes a green fluorescent protein and allows us to visualize *Cre*-expressing cell populations without inducing *Th* expression.

### Histology, Immunohistochemistry, In Situ Hybridization

Transcardiac perfusion was performed by injecting 5-10 mL of 0.1 M phosphate buffer solution into the left ventricle followed by 5 mL of 4% PFA (Fisher Scientific), both made fresh prior to perfusion. Whole brain tissue was removed and stored in 4% PFA at 4°C for 24-hours, then placed in 0.1 M PBS. 50-µm coronal and sagittal sections were obtained using a Leica VT1000S Vibratome (Leica Biosystems, Buffalo Grove, IL) and stored in 0.1M PBS at -4°C.

Tissue samples were placed in glass well plates in 0.1 M PBS for 1-min, then permeabilized by incubating in PBST (0.5% Triton X-100 in PBS) for 10-min. Samples were then blocked in 5% goat serum (Gibco, #16210-064) solution for 10-min. Tissue samples were then transferred to a well of primary antibody diluted in goat serum overnight at 4°C. After primary antibody staining, tissue was then washed three times in PBS for 5-min each. Tissue was then submerged in a dilution of secondary antibody in PBS for 1-hour shielded from light at room temp. The tissue samples were washed in PBS two times for 5-min each then added to a DAPI solution for 5-min. Tissue samples were washed again for 5-min in PBS before mounting and cover slipping.

Immunofluorescence antibody staining was performed using: chicken polyclonal TH (1:500, Aves, #TYH), rat monoclonal DAT (1:250, Millipore Sigma, #MAB369), rabbit polyclonal GFP (1:500, Invitrogen, #A6455 & #A11122), rabbit polyclonal RFP (1:500, Rockland, 600-401-379) primary antibodies. Secondary staining was performed using: Alexa Fluor 647-conjugated Affinipure Goat Anti-Chicken IgG (1:500, Invitrogen, A21449), AffiniPure Goat Anti-Rat 568 IgG (1:500, Invitrogen, A11077), and Goat Anti-Rat 488 IgG (1:500, Invitrogen, A11006), alongside DAPI (4’,6-Diamidino-2-Phenylindole, Dihydrochloride, 1:1000, Biotium, #40011) for nuclei staining. Sections were mounted using Immu-Mount (Epredia, #9990402) and imaging was performed using a Nikon C2 Confocal scope with a standard detector system and Nikon Elements Software. Adjustments to images were performed with ImageJ (Fiji) software on a separate computer system.

Fluorescent *In Situ* Hybridization (FISH) protocol was adapted from Choi et al. (2018).^55^ In brief, complementary single-stranded DNA probes were added against the following mRNAs: *Th* (GenBank accession number: NM_009377.1), *Slc6a3* (NM_010020.3), and *Calb1* (P05937). The probes were incubated with free-floating sections for ∼24-hours inside a humidified chamber at 37°C. Next, fluorescent DNA hairpins, which hybridize with initiator sequences located on the mRNA binding probes and form amplification polymers, were added for ∼24-hours at room temperature away from light. Afterward, tissues were washed with 5x SSCT at room temperature and DAPI was added for nuclei staining before mounting and cover slipping. For whole brain imaging of *Slc6a3^Cre^;Th^Flox^*cKO we utilized passive CLARITY.^36^ After confocal image acquisition and post processing, cell quantification was performed with ImageJ (Fiji) using the Cell Counter plugin to manually count cells expressing separate (singular expression) or overlapping (double expression) fluorescent cellular markers.

### Statistical Analysis

Statistical analyses were performed using GraphPad Prism (Version 10.0). All data are presented as mean ± SEM. Threshold for statistical significance was pre-defined as P < 0.05. For data sets collected over the 28-day CR protocol (Raw/Normalized daily body weights, locomotor activity, FAA activity, FED 3 wheel and nose poke data), a Mixed-Effects Model was used to analyze significance with Sidak’s multiple comparisons test. For singular comparisons between two groups (Raw/Normalized total food intake, cell counts, wheel activity, nose poking), an unpaired two-tailed Student’s t-Test with Welch’s correction was used. Pitx3-Cre strain survival was evaluated using the Log-Rank (Mantel-Cox) test. P values on all figures are denoted as follows: *≤0.05, **≤0.01, ***≤ 0.001, ***≤0.0001. Graphs were prepared using GraphPad Prism and figure layouts created using Adobe Illustrator.

## Supporting information

Control CLARITY TH staining

cKO 1 CLARITY TH staining

cKO 2 CLARITY TH staining

cKO 3 CLARITY TH staining

cKO 4 CLARITY TH staining

## Acknowledgments

We are grateful to Dr. Craig LaMunyon (CPP) for help with confocal microscopy, to Kevin Chung (CPP) for laboratory management, and to Dr. Viviana Gradinaru (Caltech) for assistance with CLARITY experiments.

## Declaration of generative AI and AI-assisted technologies in the manuscript preparation process

During the preparation of this manuscript, the authors used ChatGPT and Gemini to improve the clarity and readability of the text. After using this tool/service, the authors reviewed and edited the content as needed and take full responsibility for the content of the published article.

## Supplemental Figure Legends

**Figure S1. The impact of *Pitx3^Cre^* mediated deletion on survival and midbrain co-expression TH and DAT.** A) Survival plot of *Pitx3^Cre^ cKO* (red) mice with conditionally deleted TH compared to control (black) mice. B) Confocal imaging of *in situ* hybridization of TH (magenta) and DAT (green) in the midbrain of WT mouse tissue. Overlapping expression identifies a small subset of TH-positive, DAT-negative neurons in the midbrain (green cells). C) Residual TH-positive neurons in cKO mice visualized using passive CLARITY based immunolabeling and whole-brain map of undeleted dopamine neurons. D) Raw body weight of cKO and WT mice over period of study.

**Figure S2. AAV-RFP-TH Rescue of *Slc6a3^Cre^;Th^Flox^*cKO Mice.** A) Confocal imaging of *in situ* hybridization of TH (magenta) and RFP (green) in the midbrain of cKO rescue mouse tissue. RFP presence indicates cells with TH expression restored. B) Confocal imaging of *in situ* hybridization of TH (magenta) and GFP (green) in the midbrain of WT mouse tissue. C) Quantification of RFP+/TH+ staining in the SN of *Slc6a3^Cre^;Th^Flox^*(orange) midbrain tissue. D) Quantification of GFP+ staining in the SN of WT (gray) midbrain tissue. E) Raw body weight of WT-GFP and *Slc6a3^Cre^;Th^Flox^* cKO rescue mice over the course of the study. F) Raw food intake of WT-GFP and *Slc6a3^Cre^;Th^Flox^* cKO rescue mice over the course of the study.

**Figure S3. Characterization of Multiple Conditional TH-Flox cKO mutant lines.** A) Normalized high activity of WT (gray) and *Crhr1^Cre^;Th^Flox^* cKO (pink) mice over the course of the study; data is normalized by dividing the seconds of high activity by the total daily high activity. B) Normalized high activity of WT and *Crhr1^Cre^;Th^Flox^*cKO mice during the 3-hour premeal window throughout the course of the study; data is normalized by dividing the seconds of high activity in the premeal window by the total daily high activity. C) Raw body weight of WT and *Crhr1^Cre^;Th^Flox^*cKO mice over the course of the study. D) Normalized body weight of WT and *Crhr1^Cre^;Th^Flox^*cKO mice over the course of the study; weight was normalized by dividing the weekly weight by the Day 0 weight, which was taken prior to beginning CR. E) Raw food intake of WT and *Crhr1^Cre^;Th^Flox^* cKO mice while on AL. F) Normalized food intake of WT and *Crhr1^Cre^;Th^Flox^*cKO mice. Food intake was normalized by dividing average 24 hour AL intake by the animal’s body weight. G) Normalized high activity of WT (gray) and *Foxp2^Cre^;Th^Flox^*cKO (green) mice over the course of the study; data is normalized by dividing the seconds of high activity by the total daily high activity. H) Normalized high activity of WT and *Foxp2^Cre^;Th^Flox^* cKO mice during the 3-hour premeal window throughout the course of the study; data is normalized by dividing the seconds of high activity in the premeal window by the total daily high activity. I) Raw body weight of WT and *Foxp2^Cre^;Th^Flox^*cKO mice over the course of the study. J) Normalized body weight of WT and *Foxp2^Cre^;Th^Flox^*cKO mice over the course of the study; weight was normalized by dividing the weekly weight by the Day 0 weight, which was taken prior to beginning CR. K) Raw food intake of WT and *Foxp2^Cre^;Th^Flox^* cKO mice while on AL. L) Normalized food intake of WT and *Foxp2^Cre^;Th^Flox^* cKO mice. Food intake was normalized by dividing average 24 hour AL intake by the animal’s body weight. M) Normalized high activity of WT (gray) and *Ntsr1^Cre^;Th^Flox^*cKO (blue) mice over the course of the study; data is normalized by dividing the seconds of high activity by the total daily high activity. N) Normalized high activity of WT and *Ntsr1^Cre^;Th^Flox^* cKO mice during the 3-hour premeal window throughout the course of the study; data is normalized by dividing the seconds of high activity in the premeal window by the total daily high activity. O) Raw body weight of WT and *Ntsr1^Cre^;Th^Flox^*cKO mice over the course of the study. P) Normalized body weight of WT and *Ntsr1^Cre^;Th^Flox^*cKO mice over the course of the study; weight was normalized by dividing the weekly weight by the Day 0 weight, which was taken prior to beginning CR. Q) Raw food intake of WT and *Ntsr1^Cre^;Th^Flox^* cKO mice while on AL. R) Normalized food intake of WT and *Ntsr1^Cre^;Th^Flox^*cKO mice. Food intake was normalized by dividing average 24 hour AL intake by the animal’s body weight. S) Normalized high activity of WT (gray) and *Sox6^Cre^;Th^Flox^*cKO (purple) mice over the course of the study; data is normalized by dividing the seconds of high activity by the total daily high activity. T) Normalized high activity of WT and *Sox6^Cre^;Th^Flox^*cKO mice during the 3-hour premeal window throughout the course of the study; data is normalized by dividing the seconds of high activity in the premeal window by the total daily high activity. U) Raw body weight of WT and *Sox6^Cre^;Th^Flox^* cKO mice over the course of the study. V) Normalized body weight of WT and *Sox6^Cre^;Th^Flox^*cKO mice over the course of the study; weight was normalized by dividing the weekly weight by the Day 0 weight, which was taken prior to beginning CR. W) Raw food intake of WT and *Sox6^Cre^;Th^Flox^*cKO mice while on AL. X) Normalized food intake of WT and *Sox6^Cre^;Th^Flox^*cKO mice. Food intake was normalized by dividing average 24 hour AL intake by the animal’s body weight. Y) Normalized high activity of WT (gray) and *Slc17a6^Cre^;Th^Flox^*cKO (red) mice over the course of the study; data is normalized by dividing the seconds of high activity by the total daily high activity. Z) Normalized high activity of WT and *Slc17a6^Cre^;Th^Flox^* cKO mice during the 3-hour premeal window throughout the course of the study; data is normalized by dividing the seconds of high activity in the premeal window by the total daily high activity. AA) Raw body weight of WT and *Slc17a6^Cre^;Th^Flox^*cKO mice over the course of the study. AB) Normalized body weight of WT and *Slc17a6^Cre^;Th^Flox^*cKO mice over the course of the study; weight was normalized by dividing the weekly weight by the Day 0 weight, which was taken prior to beginning CR. AC) Raw food intake of WT and *Slc17a6^Cre^;Th^Flox^* cKO mice while on AL. AD) Normalized food intake of WT and *Slc17a6^Cre^;Th^Flox^* cKO mice. Food intake was normalized by dividing average 24-hour AL intake by the animal’s body weight.

**Figure S4. Additional characterization of physiology and timed feeding in *Calb1^Cre^;Th^Flox^* cKO mice.** A) Raw body weight of WT and *Calb1^Cre^;Th^Flox^* cKO mice over the course of the study. B) Raw food intake of WT and *Calb1^Cre^;Th^Flox^* cKO mice while on AL feeding schedule. C) High activity (s) over the course of 2-hrs post-cage cue on Day 23 of study. D) High activity (s) over the course of 2-hrs post-cage cue on Day -7 of study. E) Total high activity (s) over the course of 2-hours post-cage cue on Day 23 of study. F) Total high activity (s) over the course of 2-hours post-cage cue on Day -7 of study. G) High activity (s) on day -7 of the study. H-I) Normalized high activity on days 14 (H), and 28 (I); data is normalized by dividing the seconds of high activity in each hour window by the total daily high activity.

**Figure S5. Calb1^Cre^-mediated recombination in hypothalamic and other catecholaminergic populations.** A) TH immunostaining of hypothalamic dopamine populations (A11-A13) and locus coeruleus in control and *Calb1^Cre^;Th^Flox^*mice. Insets indicate A11 and A12 regions. B) Quantification of TH^+^ neurons demonstrating significant reductions in A11 and A12 populations, but not A13 or locus coeruleus. C) Adult *Calb1^Cre^* activity assessed in *Calb1^Cre^*;Ai6 mice following systemic delivery of 3E11 AAV9-PHP.eB-CAG-FLEX-tdTomato (one mouse shown that was representative of n=3 total). Ai6 reporter expression (ZsGreen) identifies historical recombination, whereas tdTomato labels cells with active adult Cre expression. **Figure S6. Viral rescue of TH expression in the SN fails to restore FAA in *Calb1^Cre^;Th^Flox^*mice.** A) Confocal imaging of immunofluorescence antibody staining of TH (magenta) and RFP (green) in the midbrain of *Calb1^Cre^;Th^Flox^*cKO midbrain tissue post TH-viral rescue attempt. B) Confocal imaging of immunofluorescence antibody staining of TH (magenta) and GFP (green) in the midbrain of *Calb1^Cre^;Th^Flox^*cKO midbrain GFP-control tissue. C) Quantification of GFP+ (gray) cells/section ± SEM in control midbrain tissue. D) Quantification of RFP+/TH+ (teal) co-expressing cells/section in TH-rescue midbrain tissue. E) Comparison of TH+ cells/section in GFP control (gray) and RFP-TH rescue (teal) midbrain tissue. F) Raw body weight of GFP control and RFP-TH rescue mice over the course of the study. G) Normalized body weight of GFP control and RFP-TH rescue mice over the course of the study. Weight was normalized by dividing the weekly weight by the Day 0 weight, which was taken prior to beginning CR. H) Raw food intake of GFP control and RFP-TH rescue mice during AL feeding. I) Normalized food intake of GFP control and RFP-TH rescue mice during AL feeding. Food intake was normalized by dividing average 24-hour AL intake by the animal’s body weight. J) High activity (s) of GFP control and RFP-TH rescue mice over the course of the study. K) High activity of GFP control and RFP-TH rescue mice during the 3-hour premeal window throughout the course of the study. L) Normalized high activity of GFP control and RFP-TH rescue mice during the 3-hour premeal window throughout the course of the study; data is normalized by dividing the seconds of high activity in the premeal window by the total daily high activity.

**Figure S7. Molecular and genetic distribution of *Calb1*-expressing dopaminergic subpopulations** A) Confocal imaging of immunofluorescence antibody staining of TH (magenta) and Calbindin1 (green) in the midbrain of *Calb1^Cre^;Th^Flox^* cKO midbrain tissue. B) Quantification of TH+ cells/section ± SEM in *Calb1^Cre^;Th^Flox^*cKO midbrain SN and VTA tissue. C) Confocal imaging of *in situ* hybridization staining of TH (magenta) and *Calb1* (green) mRNA in *Calb1^Cre^;Th^Flox^*cKO midbrain tissue. D) Confocal imaging of immunofluorescent expression of Cre-dependent multicolor TET-Bow labeling of *Calb1^Cre^;Th^Flox^*WT midbrain tissue. E) Confocal imaging of immunofluorescent antibody staining of TH (magenta) and Cre-dependent fluorescent expression of TdTomato (green) in a *Calb1^Cre^* reporter cross (Ai9). F) Adult *Calb1^Cre^*activity assessed in *Calb1^Cre^;*Ai6 mice following systemic administration of AAV9-PHP.eB-CAG-FLEX-tdTomato. TH (magenta), Ai6 lineage reporter (ZsGreen), and adult FLEX-tdTomato expression are shown in the ventral midbrain. G) Quantification of TH co-localization with adult FLEX-tdTomato and Ai6 reporter expression in the SN and VTA.

**Figure S8. Anticipatory food seeking behavior was unaltered in the mice lacking food anticipatory activity by deletion of TH in the calbindin neurons.** Control (gray); *Calb1^Cre^;Th^Flox^* (Brown). (A, B) Group-averaged 24 h profiles of wheel running activity, (D, E) rewarded left nose pokes, and (G,H) unrewarded right nose pokes under AL phase (A, D, G) and RF phase (B, E, H). Seven days of daily averages of each mouse with 30 min bin were generated first, then group average of *Calb1^Cre^;Th^Flox^*and control mice were calculated (mean ± S.E.M). Food availability is indicated by brown background; Light-dark cycle is shown by white and black boxes at the bottom of each profile. C, F, J): mean total revolutions of wheel running activity (FAA) and nose pokes (anticipatory food seeking) 4-h proceeding food availability. * p<0.05, ** p<0.01 (unpaired two-tailed Mann-Whitney test).

## Supplemental information titles and legends

**Document S1. Figures S1–S8 and Table S1**

**Table SI. Mouse strains used in this study.**

Videos S1-5. Videos of clarified midbrains stained with TH antibody. Video 1 shows a control brain stained while videos 2-5 show different examples of *Slc6a3^Cre^;Th^Flox^* mice.

