## Supplemental Figures and Table for "Genetic Dissociation of Circadian Prediction and Behavioral Output by a Calbindin1+ Dopamine Neuron Population"

### Supplemental Figure 1

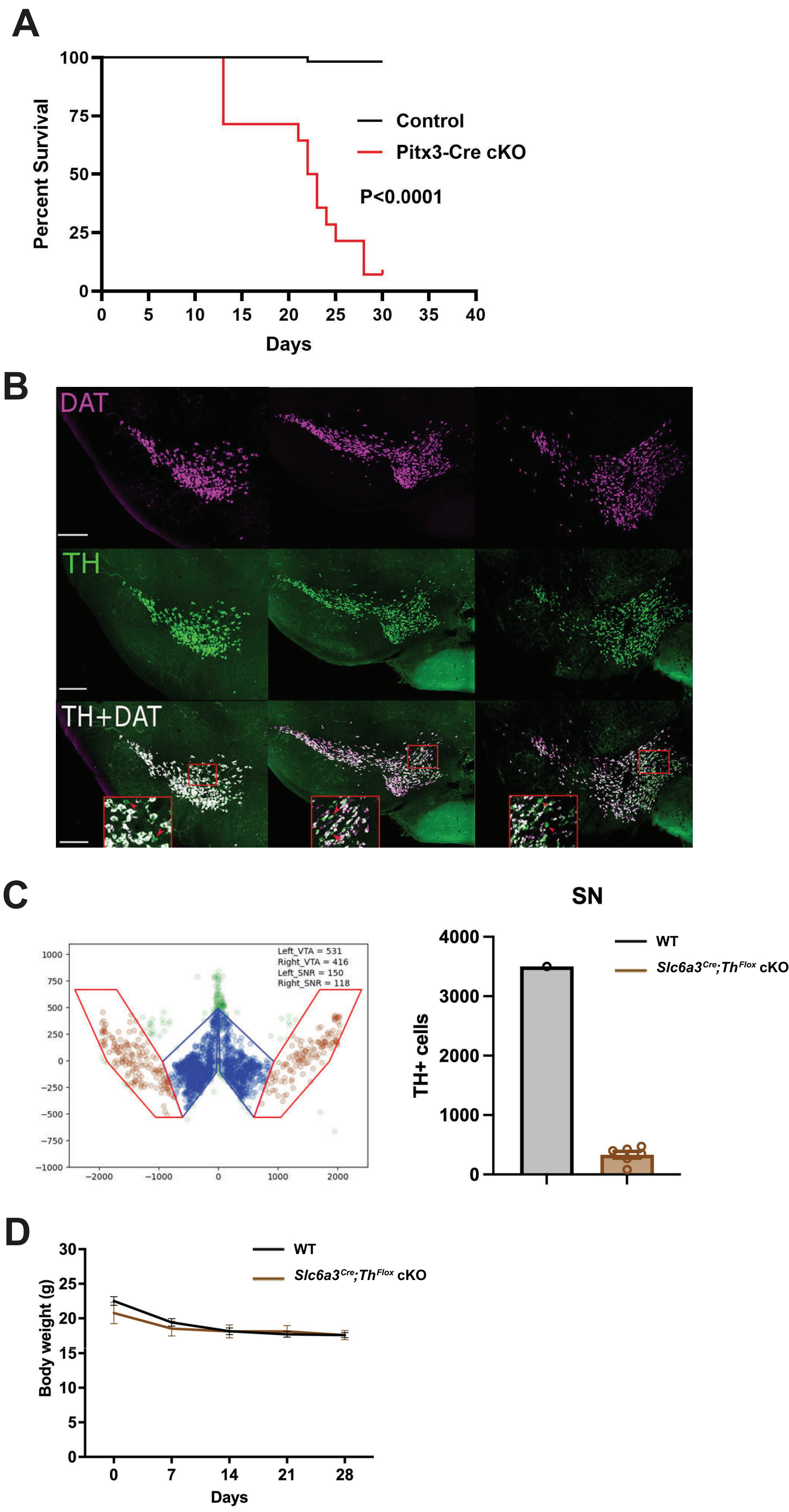

### Supplemental Figure 2

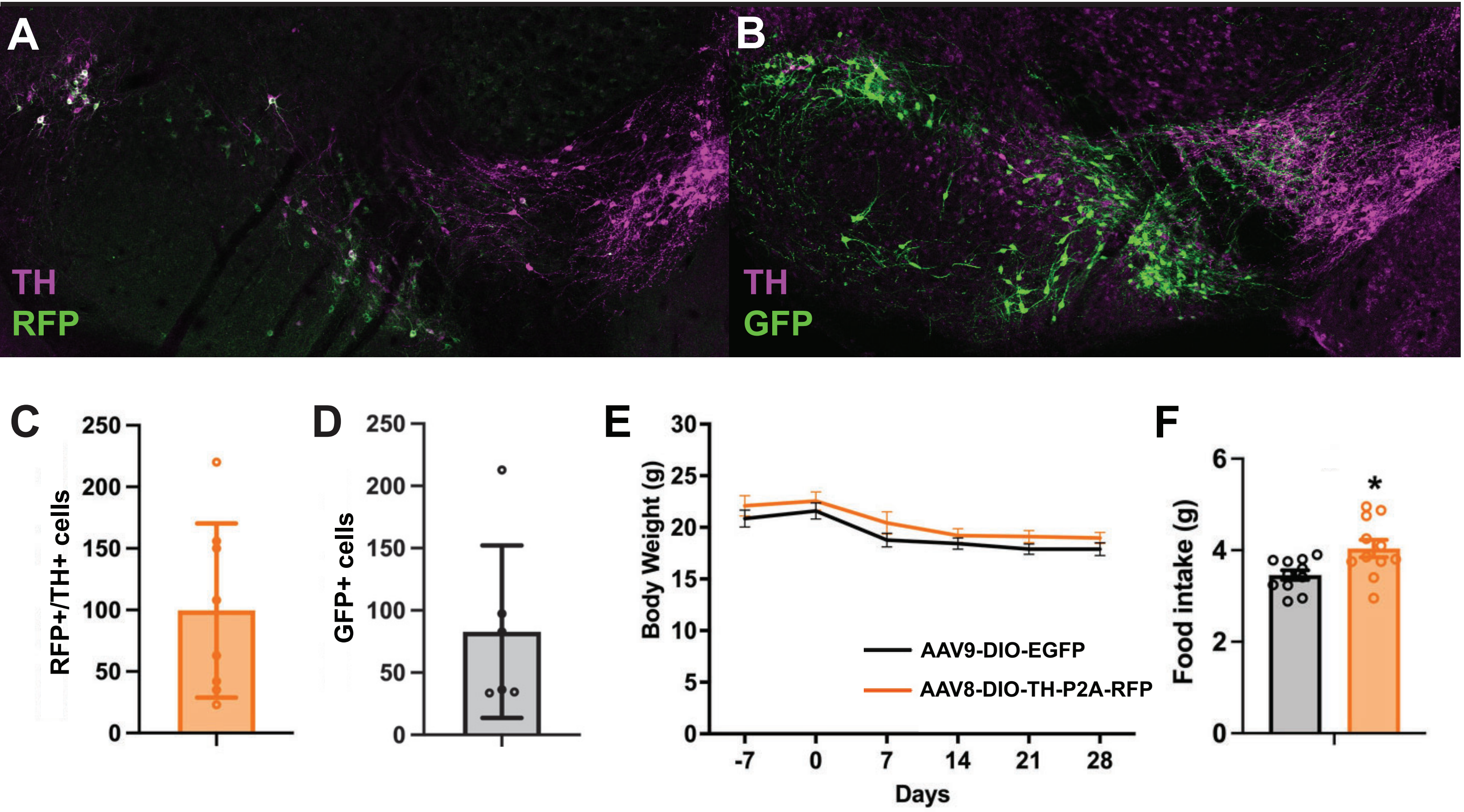

### Supplemental Figure 3

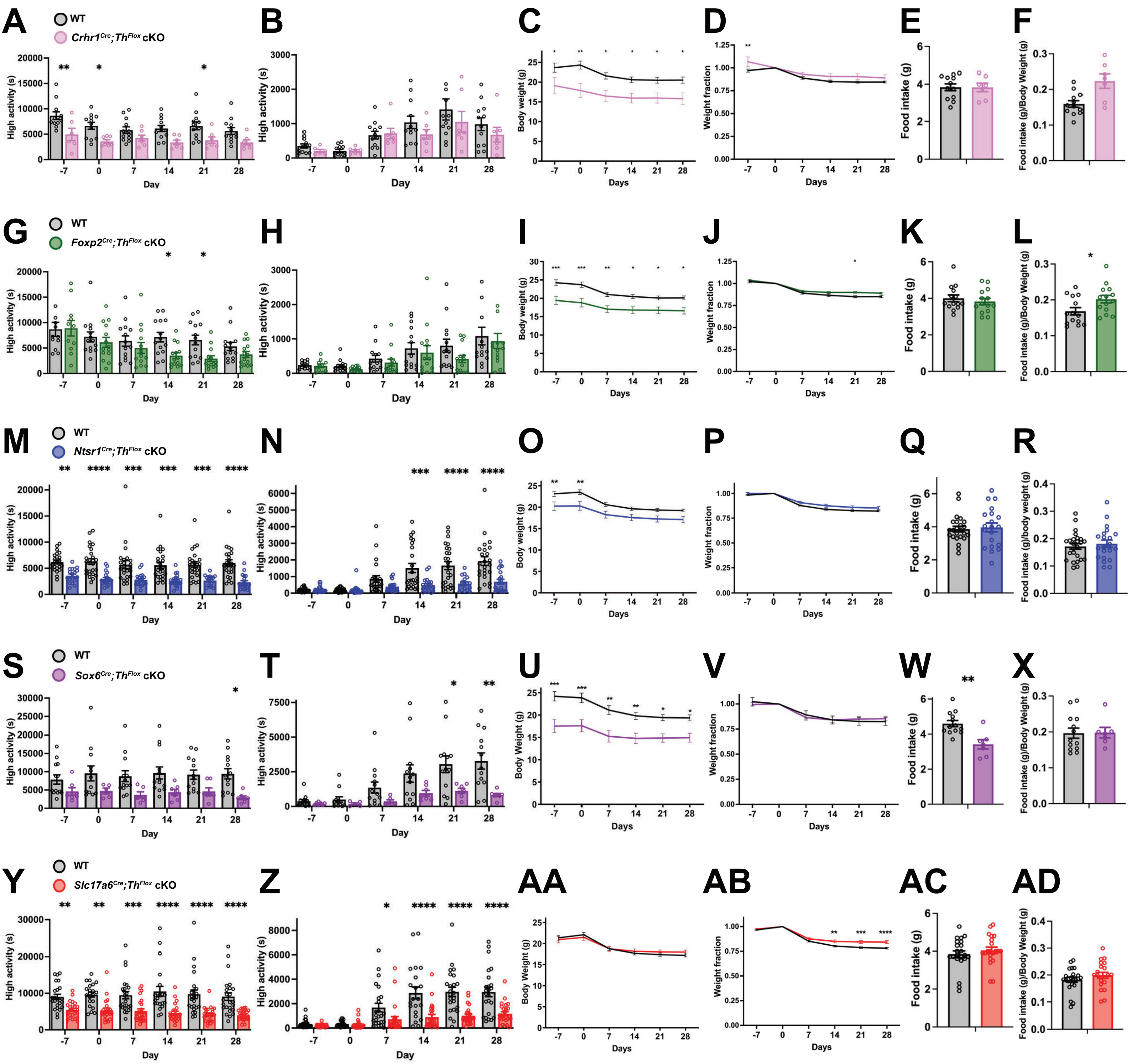

### Supplemental Figure 4

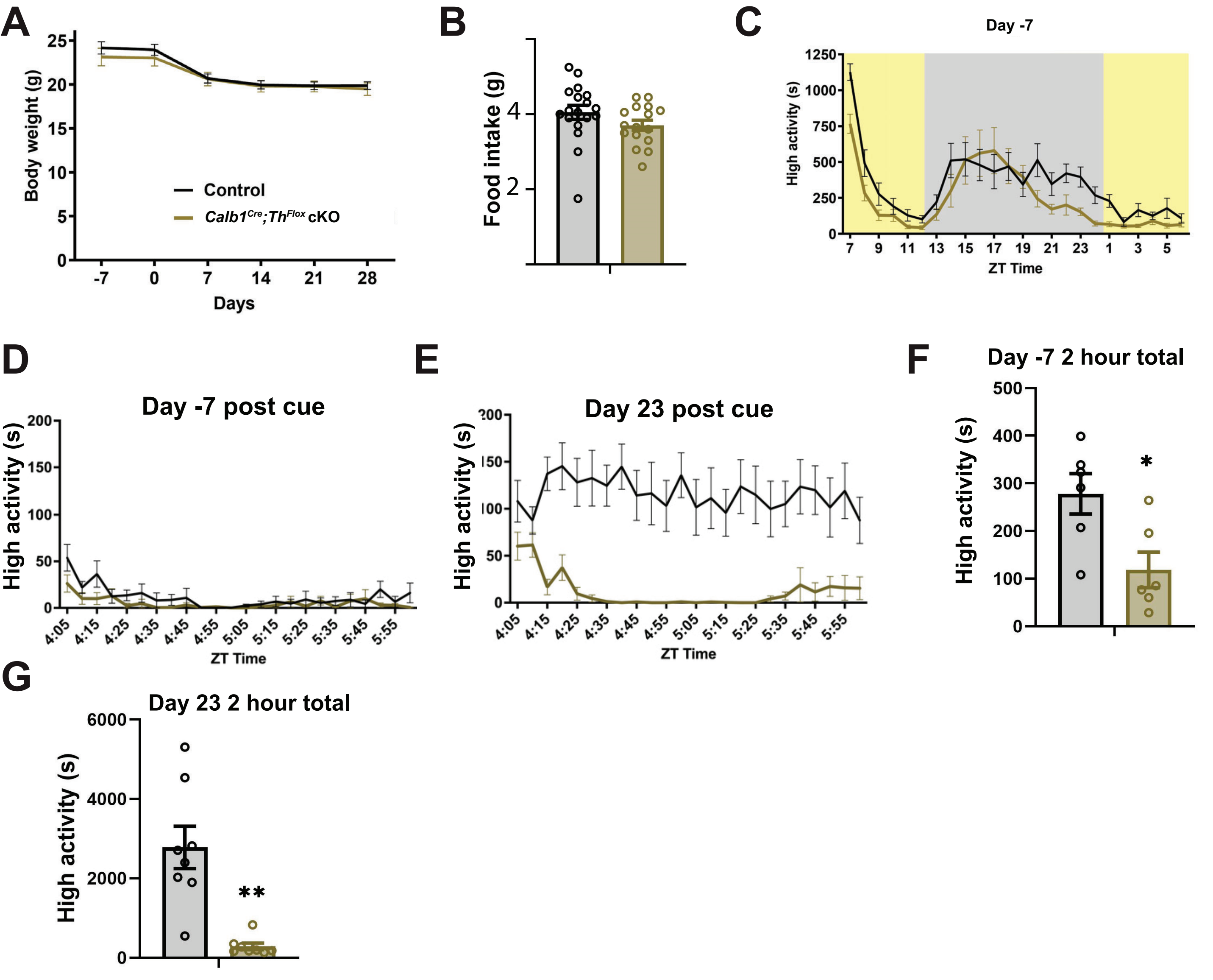

### Supplemental Figure 5

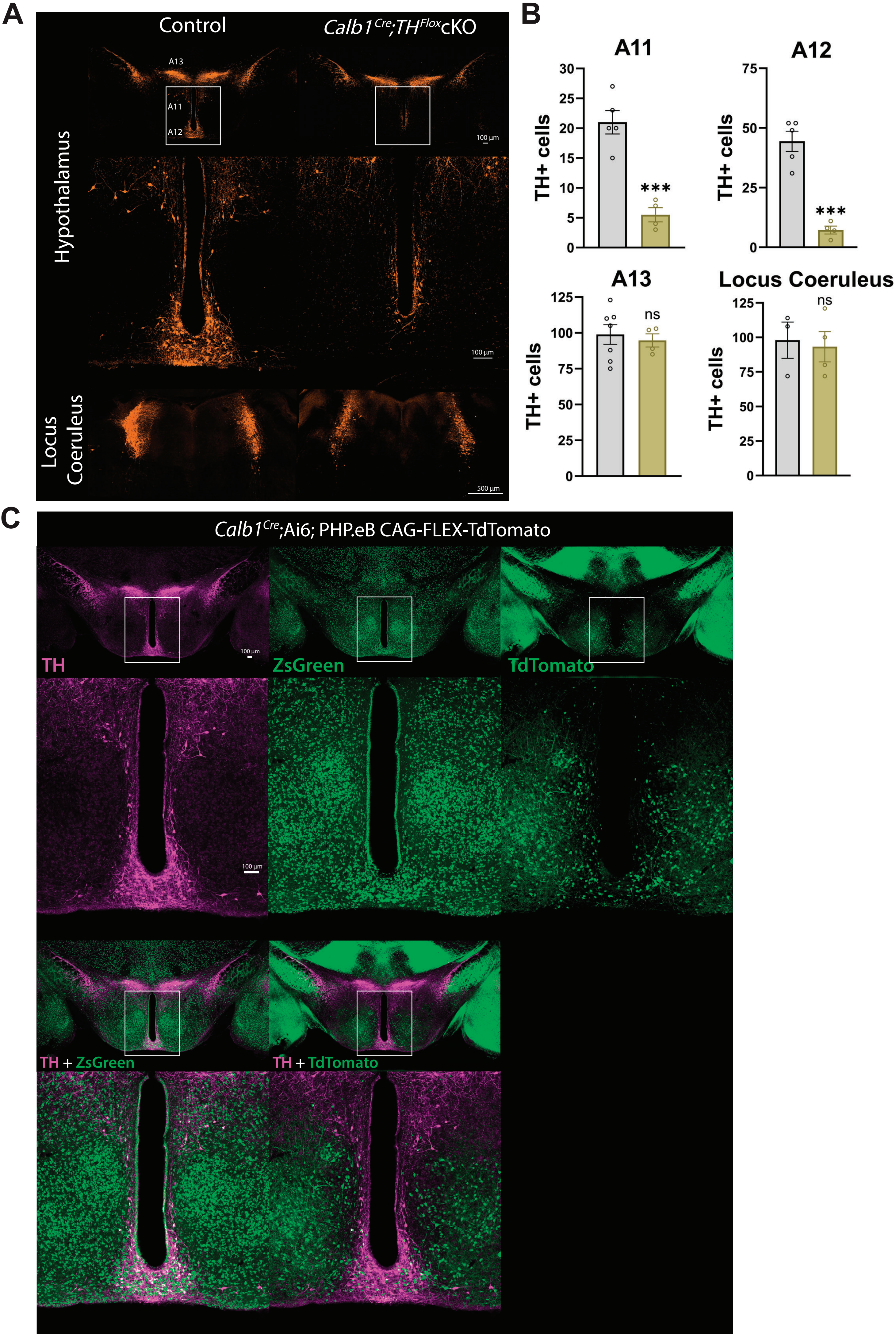

### Supplemental Figure 6

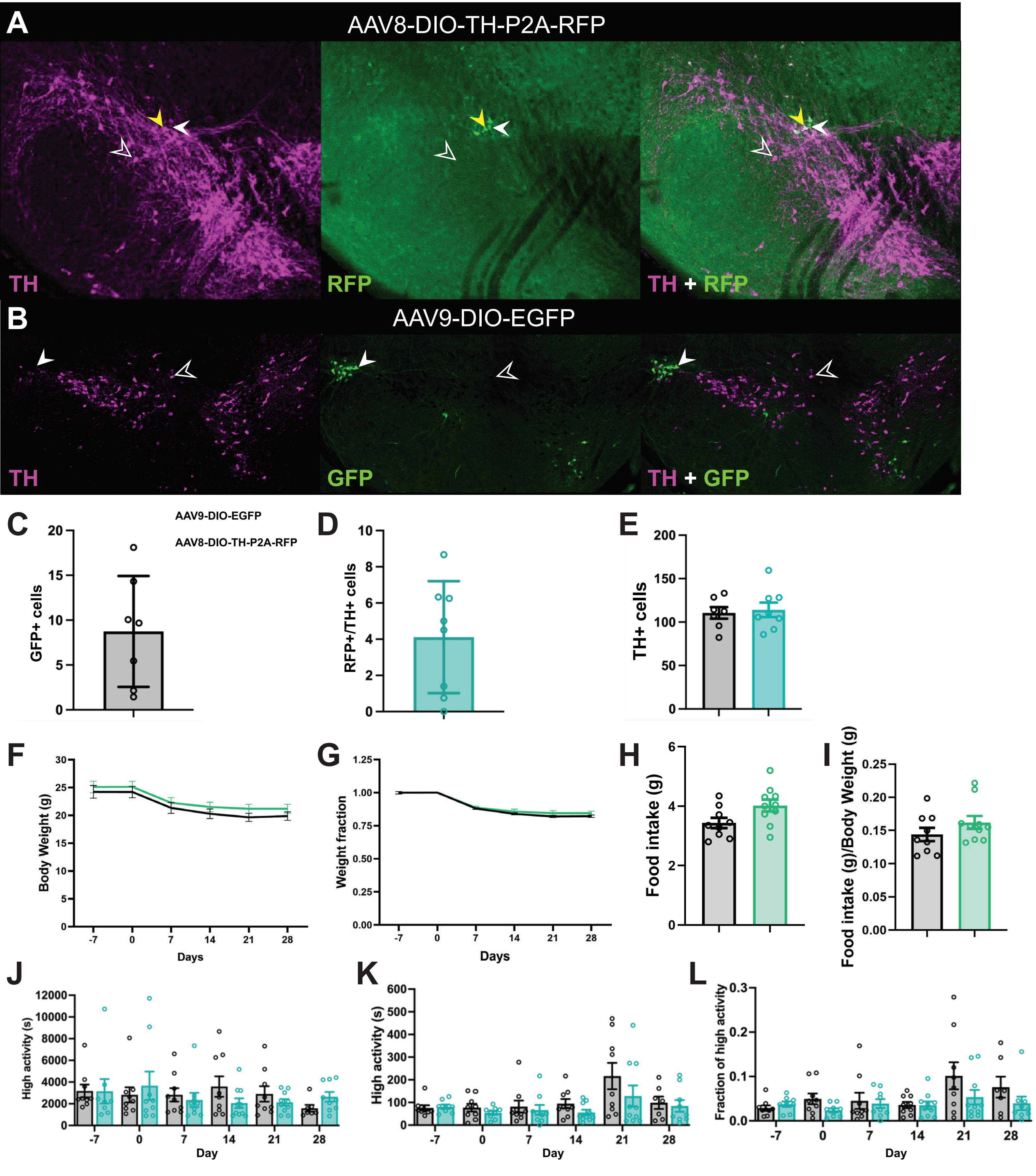

### Supplemental Figure 7

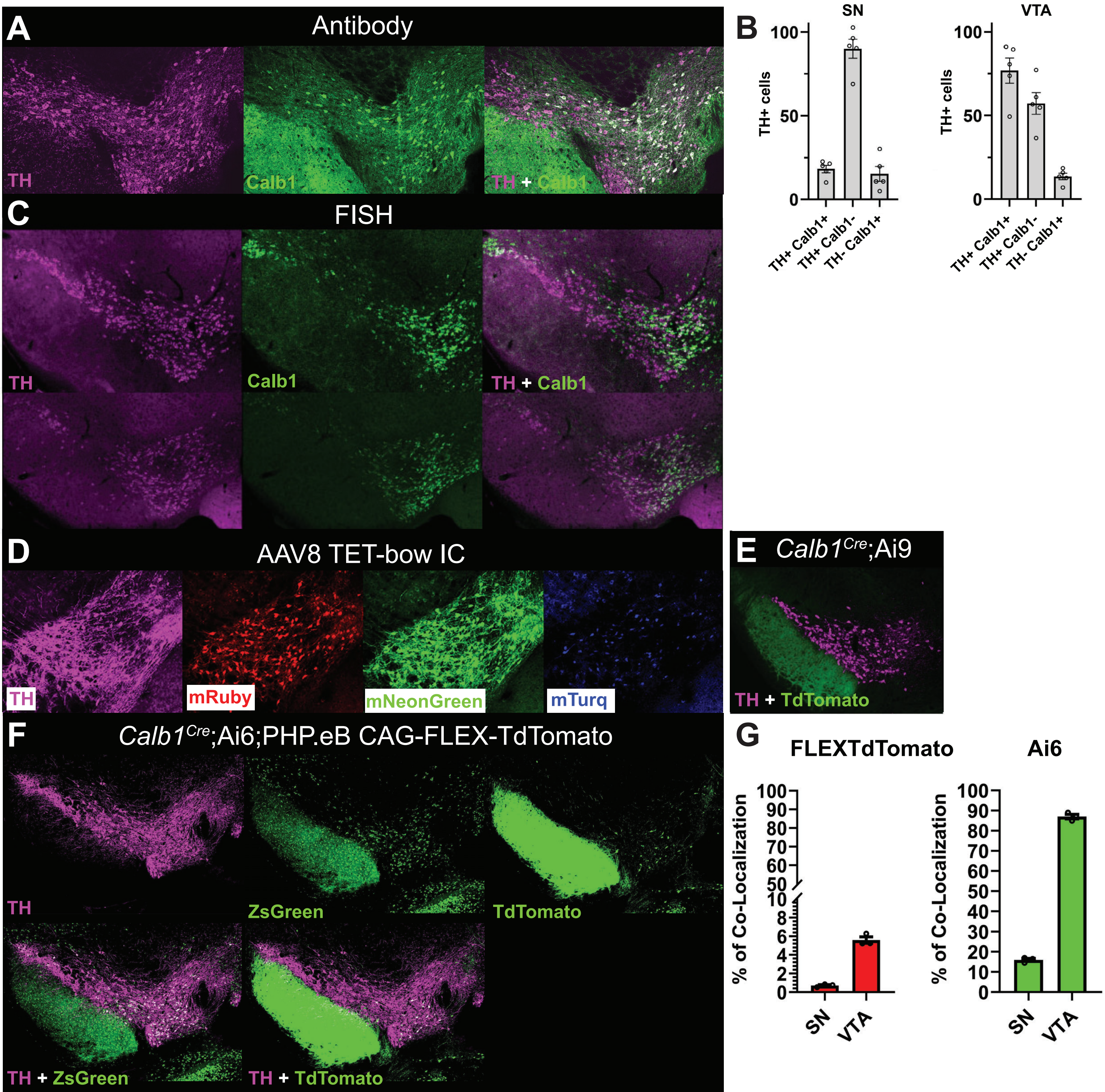

### Supplemental Figure 8

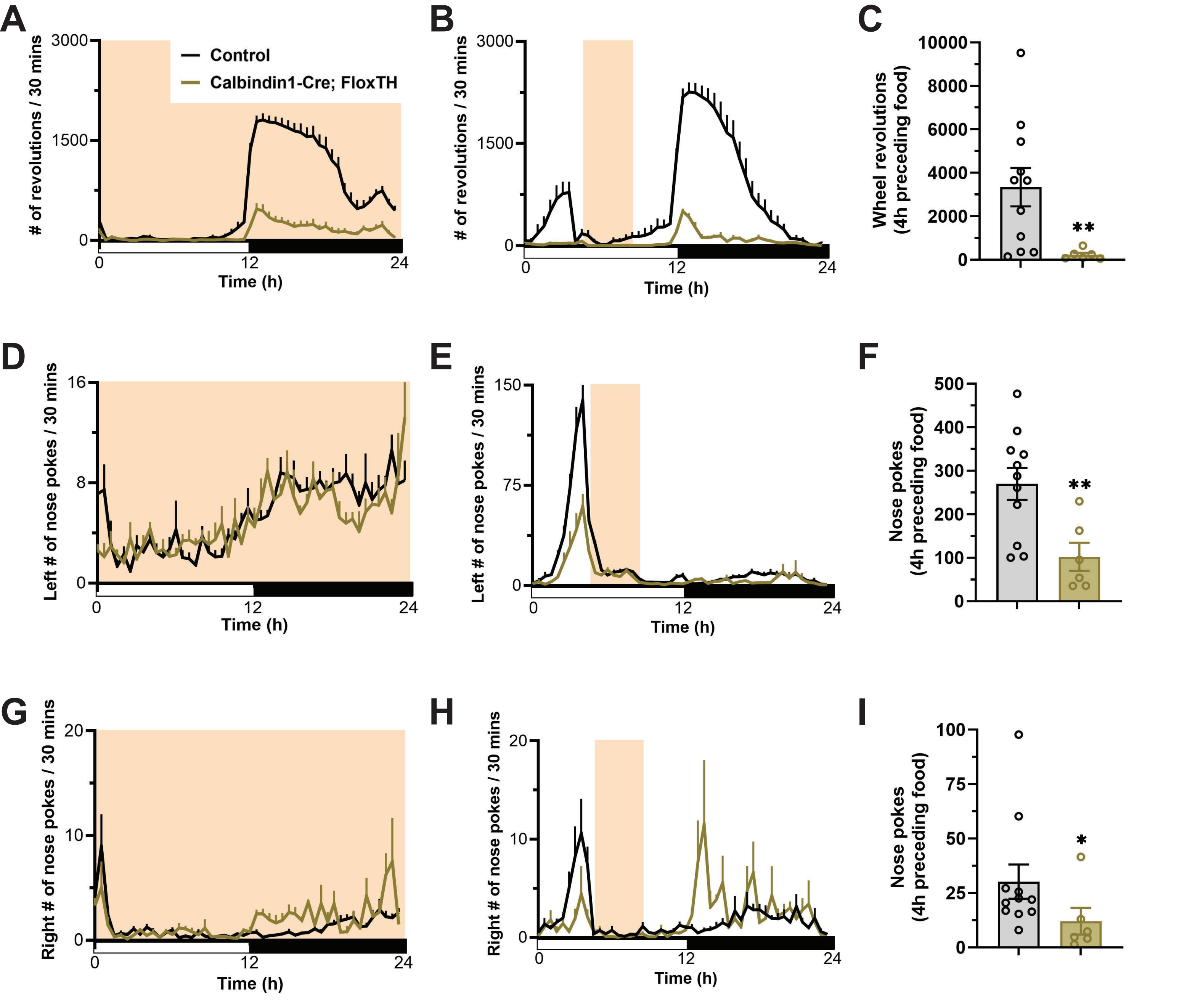

Table I. Mouse strains used in this study.

| Strain | Allele | Stock |
|---|---|---|
| *Calb1*$^{Cre}$ | *Calbindin-IRES-Cre*[45] | The Jackson Laboratory, stock #028532 |
| *Slc6a3*$^{Cre}$ | *Slc6a3-Neo-Cre*[35] | The Jackson Laboratory, stock #020080 |
| *Pitx3*$^{Cre}$ | *Pitx3-iCre*[32] | Smidt et al., 2012; Smidt Lab |
| *Crhr1*$^{Cre}$ | *Crhr1-IRES-Cre*[54] | Heymann et al., 2020; Zweifel Lab |
| *Foxp2*$^{Cre}$ | *Foxp2-IRES-Cre*[40] | The Jackson Laboratory, stock #030541 |
| *Slc17a6*$^{Cre}$ | *Slc17a6-IRES-Cre*[43] | The Jackson Laboratory, stock #016963 |
| *Ntsr1*$^{Cre}$ | *Ntsr1-ΔNeo-IRES-Cre*[41] | The Jackson Laboratory, stock #033365 |
| *Sox6*$^{Cre}$ | *Sox6-Cre*[42] | Pereira Luppi et al., 2021; Awatramani Lab |
